# Arboreal aperitifs: Space use and activity of a major songbird nest predator in a tropical Thailand forest

**DOI:** 10.1101/2021.02.09.430242

**Authors:** Anji D’souza, George Gale, Benjamin Michael Marshall, Daphawan Khamcha, Surachit Waengsothorn, Colin Thomas Strine

## Abstract

Predator-prey interactions are fundamental drivers of population dynamics, yet rarely are both predator and prey species simultaneously studied. Despite being significant, widespread avian nest predators, research on the ecology of Southeast Asian snakes in relation to birds remains scarce. The green cat snake (*Boiga cyanea*) is a primary nest predator, responsible for ≈24% of forest songbird depredation in Northeast Thailand. We explored both diurnal and nocturnal movements of 14 (5 male, 9 female) adult *B. cyanea* with radio-telemetry for an average of 68 ± 16 days per individual, between 21 October 2017 and 8 June 2019 in the dry evergreen forest of the Sakaerat Biosphere Reserve (SBR). We quantified area of space use (ha) and activity through motion variance (Ϭ_m_^2^) during the study period using dynamic Brownian bridge movement models, and linked our findings to a simultaneously-run avian nest monitoring study, initiated in 2013 within the same forest fragment. On average, movements, space use and activity differed between males and females, and between the avian nesting and non-nesting seasons. Males moved 51.37 m/day farther than females. They used areas 15.09 ha larger than females, and their activity was 3.91 Ϭ_m_^2^ higher than that of females. In general, individuals moved 50.30 m/day farther during the nesting season than the non-nesting season. The snakes used areas 9.84 ha larger during the nesting season than the non-nesting season, and their activity during the nesting season was 3.24 Ϭ_m_^2^ higher than that during the non-nesting season. All individuals were exclusively nocturnal, moving throughout the night, and often descending from higher diurnal refugia (>2 m) to forage closer to the ground after sunset. *Boiga cyanea* activity followed a similar trend to that of the recorded nest depredations at SBR. Our study links snake activity to nest depredations in SBR. Our openly-available data may yield further insight when combined with other major avian nest predator species like the congeneric invasive brown tree snake (*Boiga irregularis*) on the island of Guam.

## 1. INTRODUCTION

Declining avian populations and species richness degrade ecological processes such as seed dispersal, pollination and invertebrate population control (Sekercioglu et al., 2004). Nest predation exerts strong selection pressures on egg and nestling survival in birds; it is responsible for approximately 80% of nest failures in most species (DeGregorio et al., 2016c; Martin, 1993; Newmark and Stanley, 2011; Remeš et al., 2012). The extent and consequences of nest depredation on avian ecology have been extensively documented: it influences nest site selection, life histories and community structure (Thompson, 2007).

Reliably identifying and studying nest predators elucidates the complex predator-prey systems, allowing researchers to indirectly infer nest predator behaviour (Croston et al., 2018), while exploring habitat, temporal, and climatic characteristics that might predict nest predation risk (Sperry et al., 2009, 2008; Sperry and Weatherhead, 2009). Studies monitoring nests through cameras (Ribeiro-Silva et al., 2018) and continuous recording video systems (Pierce and Pobprasert, 2007) suggest that snakes are major, widespread avian nest predators (DeGregorio et al., 2016a; Fritts and Rodda, 1998; Robinson et al., 2005). In order to fully understand the factors that facilitate interactions between avian nests and snakes, ecologists ought to study both groups individually and simultaneously (Weatherhead and Blouin-Demers, 2004).

Research on snake-bird dynamics is limited. Snakes are cryptic and occur at low densities, making it difficult to study their foraging ecology (DeGregorio et al., 2014). The most conventional approach for quantifying the activity of cryptic taxa –like snakes, is radio-telemetry (Boback et al., 2020; Crane et al., 2020; DeGregorio et al., 2016c; Whitaker and Shine, 2003). Radio-telemetry has yielded insights into snake spatial, thermal, reproductive, and behavioural ecology (Lutterschmidt, 1994; Reinert et al., 1984; Ujvari and Korsos, 2000), while also revealing links between snake activity and resource availability, predation pressure, and temporal and environmental factors (Greene, 1997).

Few species in North America –the Eastern ratsnake *Pantherophis alleghaniensis*, the Texas ratsnake *P*. *obsoletus*, and the Corn snake *P. guttatus* (DeGregorio et al., 2016b; Sperry et al., 2008; Weatherhead and Charland, 1985)– and the introduced Brown tree snake *Boiga irregularis* on Guam (Conry, 1988; Rodda et al., 1992; Savidge, 1988) have consistent documentation regarding avian nest predation through radio-telemetry. The impact of the Brown tree snake in Guam highlights the disastrous ecological consequences of uncontrolled (introduced) snake predation on wild nesting bird populations (Fritts and Rodda, 1998; Santana-Bendix, 1994).

In Southeast Asia there is increasing evidence of snakes’ prominent role as nest predators. Khamcha *et al*. (2018) reported that snakes were responsible for 34% of predation events on 287 monitored nests. More specifically, data suggest that Green cat snake *Boiga cyanea* is the dominant snake nest predator in the seasonally wet and dry evergreen forests of Northeast Thailand (Angkaew et al., 2019; Khamcha et al., 2018; Khamcha and Gale, 2020; Pierce et al., 2020). Despite the evidence of Southeast Asian snakes– particularly that of nocturnal, arboreal species (Donald et al., 2009; Pierce et al., 2020; Pierce and Pobprasert, 2013)– as nest predators, little to no research exists assessing their movement ecology.

*Boiga cyanea* (Duméril, Bibron & Duméril, 1854) is a slender, medium to long bodied nocturnal colubrid ranging throughout Southeast Asia and Indochina. Maximum body length for females is 186 cm, and 153 cm for males (Chan-ard et al., 2015; Cox et al., 1998). Current knowledge on this species derives from occurrence records, natural history notes (Bulian and Bannasan, 1999), captive husbandry, and venom studies (Mackessy, 2002).

Our study attempts to gain insight into the free-ranging and foraging ecology of this primary nest predator as a follow up to the research by Khamcha *et al*. (2018) and Angkaew *et al*. (2019) within the same study site. We quantified adult *B. cyanea* space use and movement patterns across the avian nesting and non-nesting seasons within the Sakaerat Biosphere Reserve (SBR), Thailand. We hypothesized that space use and activity would vary between the nesting and non-nesting seasons. We predicted that (1) *B. cyanea* activity would be higher during the avian nesting season at SBR (Khamcha et al., 2018), and (2) *B. cyanea* space use would be larger during the avian non-nesting season.

## 2. MATERIAL AND METHODS

### 2.1. Study area

We conducted our research between 21 October 2017 and 8 June 2019 within the dry evergreen forests of the core area of the Sakaerat Biosphere Reserve (SBR; UNESCO-MAB Biosphere Reserve; Fig. 1), Nakhon Ratchasima Province, Thailand (14.44 – 14.55°N, 101.88 – 101.95°E). The core area of the SBR covers approximately 5,700 ha of protected area between 280 m and 762 m elevations. The core area encompasses a matrix of dry evergreen forest, open dry dipterocarp forest, transitional mixed deciduous vegetation, and operational buildings of the field research station that occupy less than 2% of the core area. *Hopea ferrea*, *Hopea odorata* and *Hydnocarpus ilicifolia* dominate the dry evergreen forest, while *Shorea obtusa*, *Shorea siamensis*, *Dipterocarpus intricatus* and *Gardenia sootepensis* are predominant characters of the dry dipterocarp forest. Marshall *et al*. (2020) summarized the annual seasonal weather patterns within the core area of SBR between 2012 and 2018 using a cluster analysis, as follows: the hot season (16 March to 30 September) averaging 33.8 ± 2.8 °C and 2.5 ± 7.9 mm/day rainfall; the wet season (1 October to 31 December) 29.9 ± 2.2 °C and 5.9 ± 11.1 mm/day rainfall; and the dry season (1 January to 15 March) 29.0 ± 3.5 °C and 0.2 ± 0.8 mm/day rainfall. However, we use Khamcha *et al*.’s (2018) definitions of wet (May to October) and dry (November to April) seasons for comparison, as they seem more biologically significant to the birds’ activity season at SBR.

**Fig. 1.**
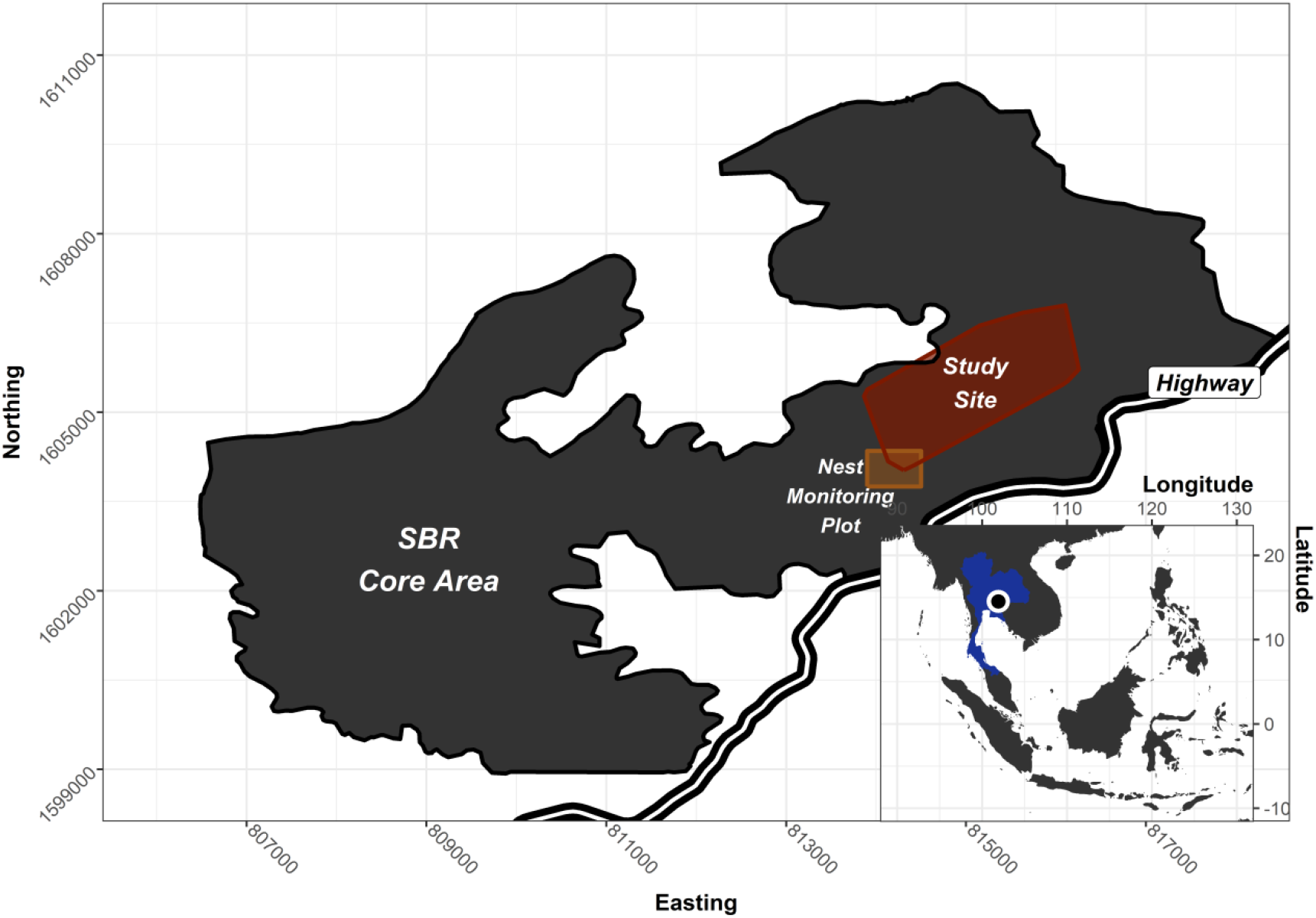
Study location. The main map shows the Sakaerat Biosphere Reserve core area. The radio-tracking study site is highlighted in red. The 36-ha nest monitoring plot is highlighted in orange. The main map is projected using UTM zone 47N, with scales in meters and north orientation. The inset map shows the study location relative to Southeast Asia, with scales in degrees.

### 2.2. Study sample

We used manual radio-telemetry to investigate *Boiga cyanea* movement patterns and space use. We obtained individuals for our research through opportunistic captures, sighting notifications from other researchers, and 904 hours of targeted nocturnal surveys in the dry evergreen forest. We measured total body lengths and mass for all captured individuals and determined their sex with cloacal probing (Schaefer, 1934). Upon collecting morphometric data, we assessed each individual’s suitability for radio-telemetry. The selection criteria required the individuals to be adults large enough to sustain an implantation surgery and accommodate a 1.8g VHF radio-transmitter (Model BD-2T or BD-2, Holohil Systems Incorporated, Ontario, Canada) in their coelomic cavity. We ensured that the radio-transmitter’s mass was less than 3% of that of the snakes. We only chose to surgically implant adult *B. cyanea* that were found within the dry evergreen forests of the core area of SBR. We also excluded adults that were excessively slender (i.e., visible neural arch or <9 mm girth at 75% snout-vent length), gravely injured or heavily gravid from being a part of the study sample.

A qualified wildlife veterinarian from Nakhon Ratchasima Zoo performed all surgeries, in accordance with Thai law. We administered isoflurane inhalation anesthetic to the snakes undergoing surgery, and followed a modified surgical technique described by Reinert and Cundall (1982). We released the implanted individuals as close as possible (≤ 20 m) to their capture location within 12 hours post-surgery to allow for recovery and thermoregulation.

### 2.3. Radio-telemetry

The BD-2T and BD-2 model Holohil radio-transmitters ideally provide a 500 – 1000 m signal range (Holohil Systems Incorporated, Ontario, Canada). However, the topography at SBR typically reduced signal range to 150 – 250 m. We located all individuals once each day during daylight hours between 21 October 2017 and 8 June 2019. We also radio-tracked the snakes during night hours after sunset but before sunrise from 14 March to 3 September, 2018, and from 15 January to 8 June, 2019. We defined fixes as the number of times we located an individual, regardless of whether it had moved or not. We defined relocations as the number of times we located an individual in a different location from its previous one.

During night-time tracks, we located each individual between one to three times at approximately four-hour intervals, every alternate night to assess the nocturnal activity and movement patterns. Exceptional circumstances during which we failed to collect data included: equipment malfunction, staff unavailability, adverse weather conditions, individuals awaiting transmitter replacement at the research facility, and inability to detect radio-signal for extended periods of time despite intensive search effort.

We estimated locations via the homing method (Amelon et al., 2009) attempting visual observations whenever feasible, primarily at night, in order to confirm the accuracy of the telemetered location. We aimed to minimize disturbance by moving discreetly through their immediate surroundings and by limiting the amount of time spent in their vicinity after pinpointing animals. We located the snakes’ daytime refugia during diurnal locations, while we assessed their activity during nocturnal fixes.

For every fix (including non-moves), we recorded the snake’s location with hand-held global positioning system (GPS) units (Garmin GPSMAP 64s, Garmin Ltd., United States) using the Universal Transverse Mercator (UTM; 47N World Geodetic System 84) coordinate reference system. We identified the snake’s position –whether arboreal (> 0 m) or terrestrial (≤ 0 m), and measured its height off the ground (m) with a measuring tape or rangefinder (Nikon Forestry Pro II, Nikon Inc., United States) for each visual observation. We documented its behaviour –whether stationary or moving, through radio-signal patterns (i.e. marked fluctuations in signal strength) or visual confirmations.

### 2.4. Nest monitoring

To explore the relationship between avian nest depredation and *B. cyanea* movement, we were granted access to nest predation records documented at the Sakaerat Biosphere Reserve by researchers from King Mongkut’s University of Technology Thonburi (KMUTT), Thailand. The nest predation data were collected within a 36-ha, permanent, nest monitoring plot in the dry evergreen forest (Khamcha et al., 2018; Khamcha and Gale, 2020; Somsiri et al., 2020) via continuous-recording video systems adapted from Pierce and Pobprasert (2007). The video systems were active throughout the nesting seasons between 2013 and 2019, monitoring 12 species of forest birds’ nests.

The dataset presents the bird species with nests depredated by *B. cyanea*, the nest heights, the date the nests were found and filmed, and the date, time and geographic coordinates of the predation events. We defined the avian nesting season in the SBR as the period between the date of discovery of the first avian nest and the date of the last nest depredation or nest abandonment. In 2018, the avian nesting season lasted from 4 February until 31 July. In 2019, the avian nesting season lasted from 4 March until 5 August. When considering the overall study period, we defined the avian nesting season as that spanning between 4 February and 5 August.

### 2.5. Software and data

We used *R v.4.0.3* (R Core Team, 2020) and *R Studio v.1.3.1093* (R Studio Team, 2020) as the front-end for data manipulation, analyses, and visualization. For data manipulation, we used packages: *dplyr v.0.8.5* (Wickham et al., 2020) to prepare our data for analysis, *lubridate v.1.7.8* (Grolemund and Wickham, 2011) to work with date and time formats, *tidybayes v.2.1.1* (Kay, 2020) to compose data for Bayesian methods, *reshape2 v.1.4.4* (Wickham, 2007) to transform data into desired formats, *raster v.3.1.5* (Hijmans et al., 2020) to read and manipulate spatial data, *rgeos v.0.5.3* (Bivand et al., 2020) to read and manipulate geographic data.

For data analyses, we used packages: *move v.4.0.0* (Kranstauber et al., 2020) to analyze animal movement data, *brms v.2.14.0* (Bürkner, 2017) to assess Bayesian generalized (non)linear multivariate multi-level models, *performance v.0.4.7* (Lüdecke et al., 2020) to assess the quality of regression models, *bestNormalize v.1.5.0* (Peterson and Cavanaugh, 2020) to find the best normalizing transformation, *overlap v.0.3.3* (Ridout and Linkie, 2009) to quantify overlap between species’ activity periods, *arm v.1.11.1* (Gelman et al., 2020) to analyze regression models, *wiqid v.0.3.0* (Meredith et al., 2020) to estimate maximum likelihood and wildlife population parameters, *adehabitatHR v.0.4.18* (Calenge, 2006) to assess habitat selection, *recurse v.1.1.2* (Bracis et al., 2018) to analyze animal trajectory data, *cluster v.2.1.0* (Maechler et al., 2019) to group and analyze data based on similarities, *MASS v.7.3.53* (Venables and Ripley, 2002) to test for significance using Pearson’s Chi-squared test, *astroFns v.4.1.0* (Harris, 2012) to convert hours to radians.

For data visualization, we used packages: *ggplot2 v.3.3.0* (Wickham, 2016), *ggpubr v.0.4.0* (Kassambara, 2020), *scales v.1.1.1* (Wickham and Seidel, 2020), *scico v.1.2.0* (Pedersen and Crameri, 2020), *ggspatial v.1.1.3* (Dunnington and Thorne, 2020), *gtable v.0.3.0* (Wickham and Pedersen, 2019), *cowplot v.1.0.0* (Wilke, 2020a), *bayesplot v.1.7.2* (Gabry and Mahr, 2020), *ggridges v.0.5.2* (Wilke, 2020b), *ggtext v.0.1.0* (Wilke, 2020c), *plotrix v.3.7.8* (Lemon et al., 2020).

Movement, activity, and dBBMM outputs and R scripts are available online at Open Science Framework (https://osf.io/6yrbg/). Movement data were additionally uploaded on Movebank (Movebank ID: 1418023557).

### 2.6. Analyses

We used the *dplyr* package (Wickham et al., 2020) to summarize data into means and their standard errors, or medians and their interquartile range (IQR) when data were non-normal or had major outliers. We used a Bayesian test of difference with Markov chain Monte Carlo (MCMC) simulations for inference on posterior distributions between groups as opposed to traditional frequentist t-tests and non-parametric tests as our sample sizes were small. We report their 95% Bayesian credible intervals (BCrI) using the Highest Density Interval (HDI) method, and their point estimates as the true difference between group means. We checked for MCMC convergence by graphically assessing their trace plots. We report a Pearson’s Chi-Squared test to assess significance (p = 0.05) between moving and stationary behaviours during the nesting and non-nesting seasons.

We estimated horizontal movements by calculating the mean daily displacement (MDD) for each individual. We defined MDD as the ratio between the sum of the Euclidean distances between consecutive fixes and the total number of days the individual was radio-tracked. We estimated space use during the study period and movement patterns using dynamic Brownian bridge movement models (dBBMM) with the *move* package (Kranstauber et al., 2020). The dynamic Brownian bridge movement models quantify an individual’s occurrence distribution based on its movement path during the study period (Kranstauber et al., 2012). This method originally analyzed GPS telemetry data on mammals and birds (Kranstauber et al., 2012). However, more recently VHF telemetry data applications on reptiles have become apparent (Knierim et al., 2019; Silva et al., 2018; Smith et al., 2020). Unlike traditional space use estimators, such as minimum convex polygons (MCP) and kernel density estimators (KDE), dBBMMs simultaneously account for spatial and temporal autocorrelation, GPS uncertainty around each location, and are more robust to irregular sampling intervals.

Additionally, dBBMMs provide estimates for an animal’s mobility –referred to as the Brownian motion variance (*Ϭ²_m_*). This metric describes heterogeneous movement behaviour or movement diffusion along an individual’s movement trajectory (Kranstauber et al., 2012), based on user-defined, biologically significant moving window and margin sizes. The moving windows and margins help estimate motion variance for subsections of the movement trajectory, thus facilitating detection of gradual and sudden changes in movement behaviour. After testing dBBMMs with larger window and margin sizes (Tables: A.1, A.2, A.3), we chose to set the window size to 9 data points to detect variations in behavioural states between 3-day periods –approximately the average time an individual would spend stationary. We set the margin size to 3 data points to detect variations between active and inactive behaviours; then specified the telemetry error associated with each data point as the mean GPS accuracy of all telemetered locations. The 90%, 95% and 99% dBBMM isopleth contours delineate snake occurrence distributions during the study period (example: Fig. A.2, Fig. A.3).

We ran a Bayesian regressive model with *brms* package (Bürkner, 2017) to assess the seasonal and sex impacts on motion variance. We opted for a Bayesian approach because the assumptions for the normality of residuals are relaxed and the estimates are more conservative. We used the *bestNormalize* package (Peterson and Cavanaugh, 2020) to shift motion variance values to a Gaussian distribution. To account for the autocorrelated structure of motion variance data, we used a third order autoregressive term (matching our selected margin size) in *brms*. We used *motion variance* as the response variable, *season* (avian nesting and non-nesting) and *sex* (male and female) as the population variables, and *individual ID* as a group effect [*motion variance* ~ *season* + *sex* + (1|*ID*) + *ar*(*p* = *3*)]. We used the package’s default priors (Student t distribution: df = 3, mu = 0, sigma = 2.5) as we had no reliable prior information to base our motion variance on for this species. We ran 5 chains with 5000 iterations with 1000 iterations of warmup and determined model convergence with trace plots and when rHat neared one (Gelman and Rubin, 1992). We coded female *B. cyanea* during the avian nesting season as the intercept, and the non-nesting season and males as the other two coefficients. We used the *bayesplot* package (Gabry and Mahr, 2020) as a visual diagnosis for autocorrelation of all model variables, and for posterior predictive checking of observed data compared to simulated data. We used the *performance* package (Lüdecke et al., 2020) to derive the R-squared (*R*^2^) regression metric to estimate the proportion of variation explained by the predictor variables.

## 3. RESULTS

### 3.1. Radio-telemetry

We radio-tracked a total of 14 adult *Boiga cyanea* –5 males and 9 females. We gathered a total of 1317 fixes and 640 relocations. We recorded 907 fixes during daylight hours, and 410 fixes at night. We recorded 780 fixes during the avian nesting season (diurnal: 480; nocturnal: 300), and 537 fixes during the non-nesting season (diurnal: 427; nocturnal: 110).

Tracking durations varied considerably among individuals. Most heterogeneity in duration resulted from individual loss: premature transmitter failures (n = 8), inexplicable deaths (n = 3), predations (n = 2), and inaccessible recapture locations (n = 1). Of the 14 radio-tracked *B. cyanea*, we were able to successfully recapture and re-implant only 2 individuals –F01 and M04.

We only report movement and dBBMM summaries for 12 individuals, omitting F06 and F07 with 18 and 7 day tracking durations respectively (Table 1). We were unable to derive dBBMM outputs for both individuals given our window (9) and margin sizes (3).

**Table 1.** Tracking summaries of radio-tracked *Boiga cyanea* between 21 October 2017 and 8 June 2019.

| ID | SVL (mm) | Start date<br>(d/m/y) | End date<br>(d/m/y) | Days tracked | Fixes | Relocations | Time lag (hr) | Revisit frequency<br>(days) | Time stationary<br>(days) |
| --- | --- | --- | --- | --- | --- | --- | --- | --- | --- |
| F01 | 914 | 21/10/2017 | 17/02/2018 | 119 | 102 | 72 | 28.22 ± 2.7 | 18.80 ± 3.05 | 2.18 ± 0.30 |
| F02 | 973 | 23/10/2017 | 13/01/2018 | 82 | 79 | 40 | 25.18 ± 0.92 | 11.10 ± 0.87 | 5.75 ± 0.65 |
| F03 | 1086 | 28/12/2017 | 01/05/2018 | 124 | 146 | 69 | 20.51 ± 0.9 | 16.60 ± 1.21 | 3.04 ± 0.23 |
| M01 | 1046 | 23/03/2018 | 06/07/2018 | 105 | 134 | 59 | 18.96 ± 1.03 | 22.60 ± 1.62 | 9.59 ± 1.27 |
| M02 | 1200 | 03/03/2018 | 08/04/2018 | 36 | 48 | 25 | 18.36 ± 1.13 | 2.28 ± 0.23 | 1.60 ± 0.18 |
| M03 | 1150 | 24/04/2018 | 28/07/2018 | 95 | 129 | 83 | 17.81 ± 1.17 | 12.30 ± 2.46 | 1.69 ± 0.23 |
| F04 | 1204 | 11/08/2018 | 04/09/2018 | 24 | 44 | 28 | 13.53 ± 1.02 | 2.21 ± 1.06 | 1.60 ± 0.30 |
| M04 | 960 | 06/11/2018 | 08/06/2019 | 214 | 395 | 168 | 13.03 ± 0.5 | 28.10 ± 1.35 | 2.25 ± 0.13 |
| F05 | 1205 | 12/11/2018 | 16/01/2019 | 65 | 73 | 33 | 21.6 ± 0.98 | 11.80 ± 1.15 | 5.73 ± 0.71 |
| F06 | 1114 | 12/11/2018 | 01/12/2018 | 18 | 25 | 2 | 18.45 ± 1.31 | 0.61 ± 0.08 | 5.49 ± 0.62 |
| F07 | 1028 | 23/03/2019 | 30/03/2019 | 7 | 15 | 4 | 11.83 ± 2.29 | 1.15 ± 0.15 | 1.49 ± 0.09 |
| F08 | 952 | 07/04/2019 | 29/04/2019 | 22 | 46 | 18 | 11.55 ± 1.91 | 2.70 ± 0.00 | 2.22 ± 0.34 |
| F09 | 1050 | 27/04/2019 | 24/05/2019 | 27 | 48 | 16 | 13.9 ± 1.51 | 6.79 ± 1.14 | 3.00 ± 0.39 |
| M05 | 1174 | 19/05/2019 | 08/06/2019 | 19 | 33 | 23 | 14.59 ± 2.19 | - | 0.54 ± 0.15 |
| Females | 1058 ± 35 | 21/10/2017 | 24/05/2019 | 54 ± 15 | 64 ± 14 | 31 ± 8 | 20.57 ± 0.64 | 7.97 ± 2.29 | 3.39 ± 0.59 |
| Males | 1106 ± 45 | 23/03/2018 | 08/06/2019 | 94 ± 34 | 148 ± 65 | 72 ± 27 | 15.35 ± 0.41 | 16.32 ± 5.71 | 3.13 ± 1.64 |
| Total | 1075 ± 27 | 21/10/2017 | 08/06/2019 | 68 ± 16 | 94 ± 26 | 46 ± 12 | 17.63 ± 0.37 | 10.54 ± 2.49 | 3.29 ± 0.66 |

\***SVL:** Snout-to-Vent Length; SVL, days tracked, fixes, relocations, time lag(as seen in Fig. A1), revisit frequency and time stationary for females, males and the whole sample are reported as means ± standard error.

### 3.2. Space use

#### 3.2.1. Horizontal movements

The mean daily displacement (MDD; Table 2) for male and female snakes together was short –just over 55 m/day. Males moved significantly longer daily distances than females (Point Estimate of Difference (PED): 51.37 m/day; 95% Bayesian Credible Interval (BCrI): 5.05 – 97.9). Snakes moved significantly greater daily distances on average during the avian nesting season compared to the non-nesting season (PED: 50.30 m/day; 95% BCrI: 21.1 – 79.1). We caution that the samples may not be independent, as individuals may have influenced other individuals’ movements by increasing or reducing them.

**Table 2.** Movement summaries of radio-tracked *Boiga cyanea.*

| Snake ID | SVL (mm) | MDD (m/day)<br>Nesting | MDD<br>(m/day)<br>Non-nesting | MDD<br>(m/day) | Max. distance<br>moved (m) | Max. days<br>stationary |
| --- | --- | --- | --- | --- | --- | --- |
| F01 | 914 | 10 | 25 | 23 | 161 | 11 |
| F02 | 973 | - | 15 | 15 | 132 | 24 |
| F03 | 1086 | 52 | 9 | 38 | 260 | 13 |
| F04 | 1204 | - | 28 | 28 | 68 | 3 |
| F05 | 1205 | - | 29 | 29 | 400 | 23 |
| F08 | 952 | 62 | - | 62 | 402 | 4 |
| F09 | 1050 | 50 | - | 50 | 167 | 9 |
| M01 | 1046 | 100 | - | 100 | 385 | 33 |
| M02 | 1200 | 46 | - | 46 | 190 | 4 |
| M03 | 1150 | 115 | - | 115 | 583 | 4 |
| M04 | 960 | 87 | 20 | 50 | 245 | 11 |
| M05 | 1174 | 121 | - | 121 | 524 | 1 |
| Females | 1055 ± 44 | 43.29 ± 11.46 | 21.18 ± 3.97 | 35.08 ± 6.11 | 227.14 ± 49.74 | 12 ± 3 |
| Males | 1106 ± 45 | 93.65 ± 13.22 | 20 | 86.33 ± 15.95 | 385.4 ± 76.21 | 11 ± 6 |
| Total | 1076 ± 31 | 71.27 ± 12.2 | 21.05 ± 3.25 | 56.44 ± 10.41 | 293.08 ± 47.12 | 12 ± 3 |

\*SVL: Snout-to-Vent Length; MDD: Mean Daily displacement; Max. distance moved (m): farthest distance travelled between consecutive diurnal fixes.

#### 3.2.2. Vertical movements

We confirmed 158 height observations, of which 48 in daylight and 110 at night. During the day, all height observations were 10 m or below (range: 0 – 10). Of the 48 of 907 total daylight fixes during our study, the snakes sheltered on average in the understory (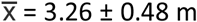; n = 31; on ground observations were not included in arboreal refugia average calculations). Proximity to the snakes and our detection ability influenced whether we could confirm daytime refugia heights, so we cannot provide reliable statistics on higher diurnal refugia. We feel confident in stating that for the remaining 859 diurnal fixes, the snakes were located at heights over 2 m above the ground.

The snakes generally foraged close to the ground at night (median height: 1.5 m; IQR: 3.5). We visually confirmed 102 night-time heights of 300 night-time fixes during the avian nesting season (Fig. 2B). The snakes moved closer to the ground during the avian nesting season (median height: 1.5 m; IQR: 3) compared to the non-nesting season (median height: 3.25; IQR: 5.42).

**Fig. 2.**
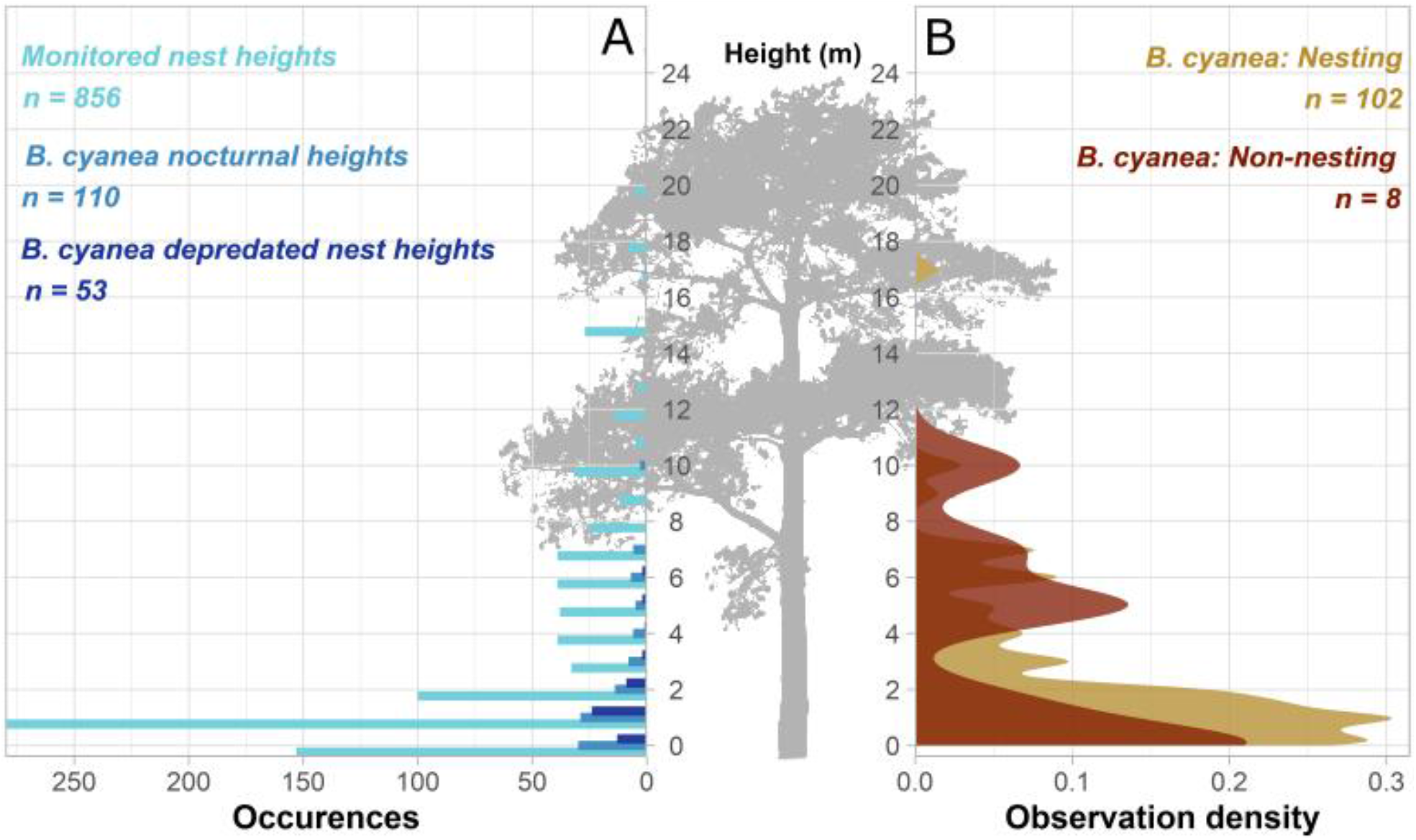
Height comparisons of *Boiga cyanea* activity and monitored avian nests at SBR. **A)** Histogram illustrating monitored nest heights (2013 – 2019), *B. cyanea* night-time movement heights (2017.10.21 – 2019.06.08), and *B. cyanea* depredated nest heights (2013 – 2019). **B)** Density plot comparing *B. cyanea* night-time movement heights during the avian nesting seasons and non-nesting seasons between 2017.10.21 and 2019.06.08.

The avian nests monitored (n = 856) at SBR between 2013 and 2019 were all below 25 m (median height: 1.5; IQR: 4.51; Fig. 2A). All records of nests depredated by *B. cyanea* (n = 53) between 2013 and 2019 were below 6 m (median height: 1 m; IQR: 1).

#### 3.2.3. Occurrence distribution

In general, males used substantially larger areas than females (PED: 15.09 ha; 95% BCrI: −3.34 – 33; Table 3). We only tracked one male during the avian non-nesting season, so we cannot infer differences in male space use between the nesting and non-nesting seasons. Females used similar areas between the nesting and non-nesting seasons (PED: 1.80 ha; 95% BCrI: −3.43 – 7.02). However, when combining both males and females, snakes used significantly larger areas during the nesting season than the non-nesting season (PED: 9.84; 95% BCrI: −0.02 – 20).

**Table 3.** Space use summaries of radio-tracked *Boiga cyanea.*

| Snake ID | Days tracked | dBBMM 90%<br>(ha) | dBBMM 95%<br>(ha) | dBBMM 99%<br>(ha) | Days<br>tracked<br>Nesting | dBBMM 95%<br>(ha)<br>Nesting | Days<br>tracked<br>Non-nesting | dBBMM 95%<br>(ha)<br>Non-nesting |
| --- | --- | --- | --- | --- | --- | --- | --- | --- |
| F01 | 119 | 1.66 | 2.13 | 3.17 | 13 | 0.32 | 105 | 2.19 |
| F02 | 82 | 0.89 | 1.48 | 2.70 | - | - | 82 | 1.48 |
| F03 | 124 | 4.09 | 6.31 | 11.40 | 86 | 7.55 | 37 | 0.09 |
| F04 | 24 | 0.56 | 0.84 | 1.45 | - | - | 24 | 0.84 |
| F05 | 65 | 1.21 | 4.45 | 12.34 | - | - | 65 | 4.45 |
| F08 | 22 | 2.52 | 3.84 | 7.20 | 22 | 3.84 | - | - |
| F09 | 27 | 1.85 | 2.70 | 4.80 | 27 | 2.70 | - | - |
| M01 | 105 | 27.48 | 37.57 | 59.78 | 105 | 37.57 | - | - |
| M02 | 36 | 1.99 | 2.85 | 4.21 | 36 | 2.85 | - | - |
| M03 | 95 | 18.35 | 24.81 | 39.55 | 95 | 24.81 | - | - |
| M04 | 214 | 3.97 | 5.29 | 8.74 | 96 | 6.3 | 117 | 2.73 |
| M05 | 19 | 15.84 | 20.32 | 29.80 | 19 | 20.32 | - | - |
| Females | 66 ± 17 | 1.82 ± 0.44 | 3.11 ± 0.72 | 6.15 ± 1.63 | 37 ± 17 | 3.60 ± 1.51 | 63 ± 15 | 1.81 ± 0.75 |
| Males | 94 ± 34 | 13.53 ± 4.73 | 18.17 ± 6.43 | 28.42 ± 10.2 | 70 ± 18 | 18.37 ± 6.33 | 117 | 2.73 |
| Total | 78 ± 17 | 6.7 ± 2.55 | 9.38 ± 3.38 | 15.43 ± 5.25 | 55 ± 13 | 11.81 ± 4.27 | 72 ± 15 | 1.96 ± 0.63 |

\***dBBMM**: Dynamic Brownian bridge movement model space use estimate; **Nesting**: avian nesting season; **Non-nesting**: avian non-nesting season.

### 3.3. Temporal activity patterns

#### 3.3.1. Seasonal activity

We use the dBBMM motion variance (Ϭ_m_^2^) as a proxy for snake foraging activity. Males generally displayed higher activity than females (PED: 3.91 Ϭ_m_^2^; 95% BCrI: −0.12 – 8.16; Table 4; Fig. 3). Because we only tracked one male during the avian non-nesting season, we cannot infer differences in male activity between the nesting and non-nesting seasons. Females did not exhibit much difference in activity between the nesting and non-nesting seasons (PED: 1.08 Ϭ_m_^2^; 95% BCrI: −0.52 – 2.73). However, in general, tracked snakes were significantly more active during the nesting season than the non-nesting season (PED: 3.24 Ϭ_m_^2^; 95% BCrI: 0.874 – 5.56).

**Table 4.** Motion variance (as seen in Fig. A.4) summaries of radio-tracked *Boiga cyanea*.

| Snake ID | Days Tracked | $\sigma_m^2 \pm SE$ | Days Nesting | $\sigma_m^2 \pm SE$ Nesting | Days Non-nesting | $\sigma_m^2 \pm SE$ Non-nesting |
| --- | --- | --- | --- | --- | --- | --- |
| F01 | 119 | 0.55 ± 0.07 | 13 | 0.15 ± 0.04 | 105 | 0.63 ± 0.07 |
| F02 | 82 | 0.35 ± 0.06 | - | - | 82 | 0.35 ± 0.06 |
| F03 | 124 | 1.62 ± 0.21 | 86 | 2.17 ± 0.27 | 37 | 0.14 ± 0.07 |
| F04 | 24 | 0.34 ± 0.10 | - | - | 24 | 0.34 ± 0.10 |
| F05 | 65 | 1.16 ± 0.37 | - | - | 65 | 1.16 ± 0.37 |
| F08 | 22 | 1.87 ± 0.50 | 22 | 1.88 ± 0.50 | - | - |
| F09 | 27 | 2.24 ± 0.44 | 27 | 2.24 ± 0.44 | - | - |
| M01 | 105 | 8.34 ± 0.91 | 105 | 8.49 ± 0.91 | - | - |
| M02 | 36 | 0.90 ± 0.19 | 36 | 0.90 ± 0.19 | - | - |
| M03 | 95 | 7.77 ± 0.77 | 95 | 7.76 ± 0.77 | - | - |
| M04 | 214 | 2.21 ± 0.22 | 96 | 3.94 ± 0.38 | 117 | 0.36 ± 0.04 |
| M05 | 19 | 6.16 ± 0.92 | 19 | 5.89 ± 1.02 | - | - |
| Females | 66 ± 17 | 1.16 ± 0.29 | 37 ± 17 | 1.61 ± 0.49 | 63 ± 15 | 0.52 ± 0.18 |
| Males | 94 ± 34 | 5.08 ± 1.50 | 70 ± 18 | 5.42 ± 1.36 | 117 | 0.36 |
| Total | 78 ± 17 | 2.79 ± 0.84 | 55 ± 13 | 3.73 ± 1 | 72 ± 15 | 0.5 ± 0.15 |

\* $\sigma_m^2 \pm SE$ : Motion variance ± standard error

**Fig. 3.**
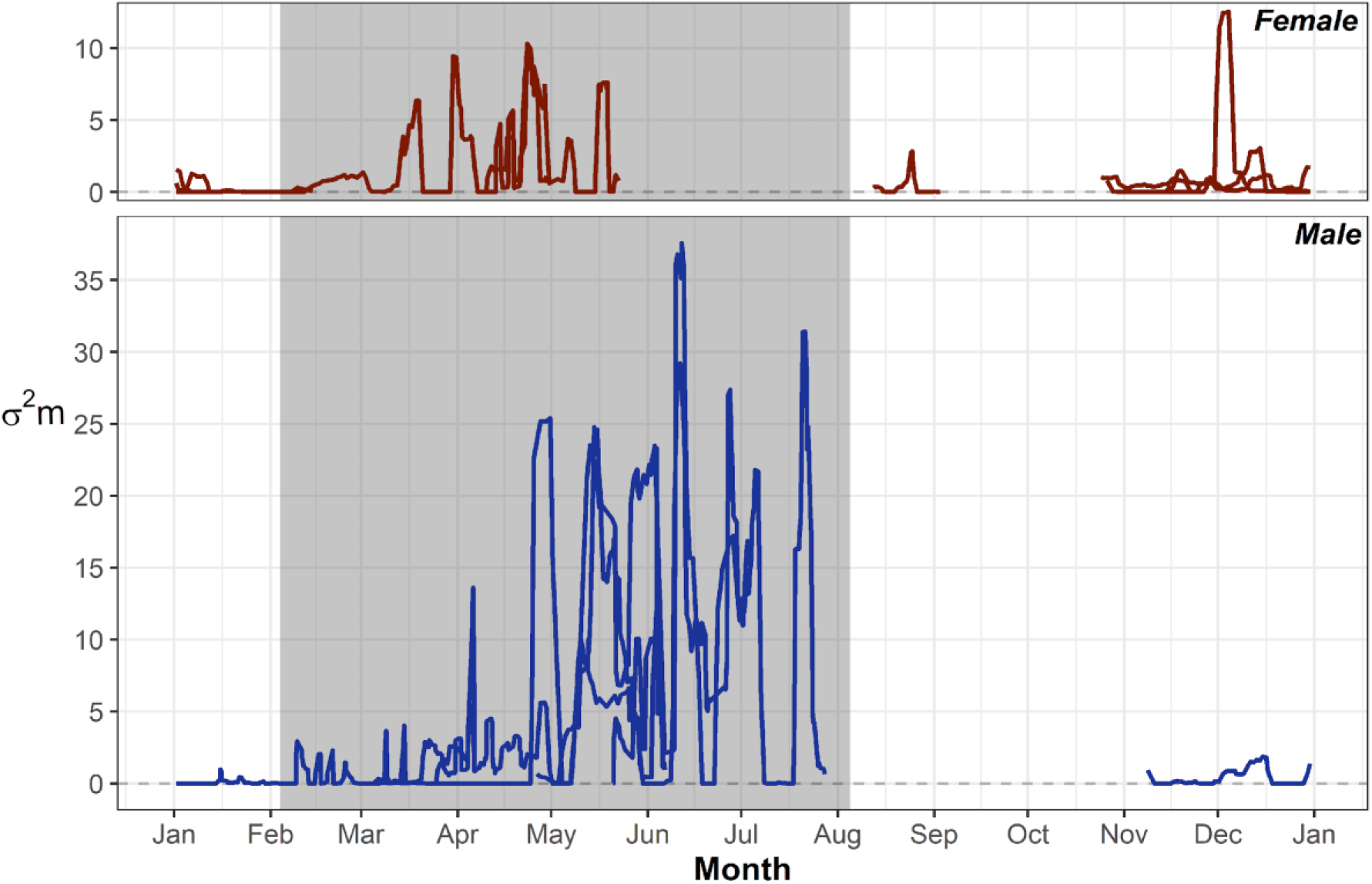
Monthly motion variances for male and female radio-tracked *B. cyanea*, with the avian nesting season (4 February – 5 August) highlighted in grey.

Our Bayesian regression model lacked evidence of autocorrelation between the model variables: the autocorrelation parameters diminish to near zero by about 25 lags. The posterior predictive check plot (Fig. A.6) suggests that our simulated data do not perfectly replicate our observed data, thus reducing model fit. Our Bayesian regression model results (Fig. 4B) further support our observations from Fig. 4A, displaying the dispersion of motion variances during the nesting and non-nesting seasons, across males and females (*R*^2^ = 0.327). The coefficients for activity during the non-nesting season (−0.5 ± 0.15; 95% BCrI: −0.35 – 0.25), and for males (−0.11 ± 0.33; 95% BCrI: −0.79 – 0.54) were both negative, suggesting reduced activity. Males exhibited more individual variation in activity, compared to females –most likely because we only radio-tracked one male during the non-nesting season. The large overlaps between coefficients suggest that season and sex are not the only factors affecting snake activity.

**Fig. 4.**
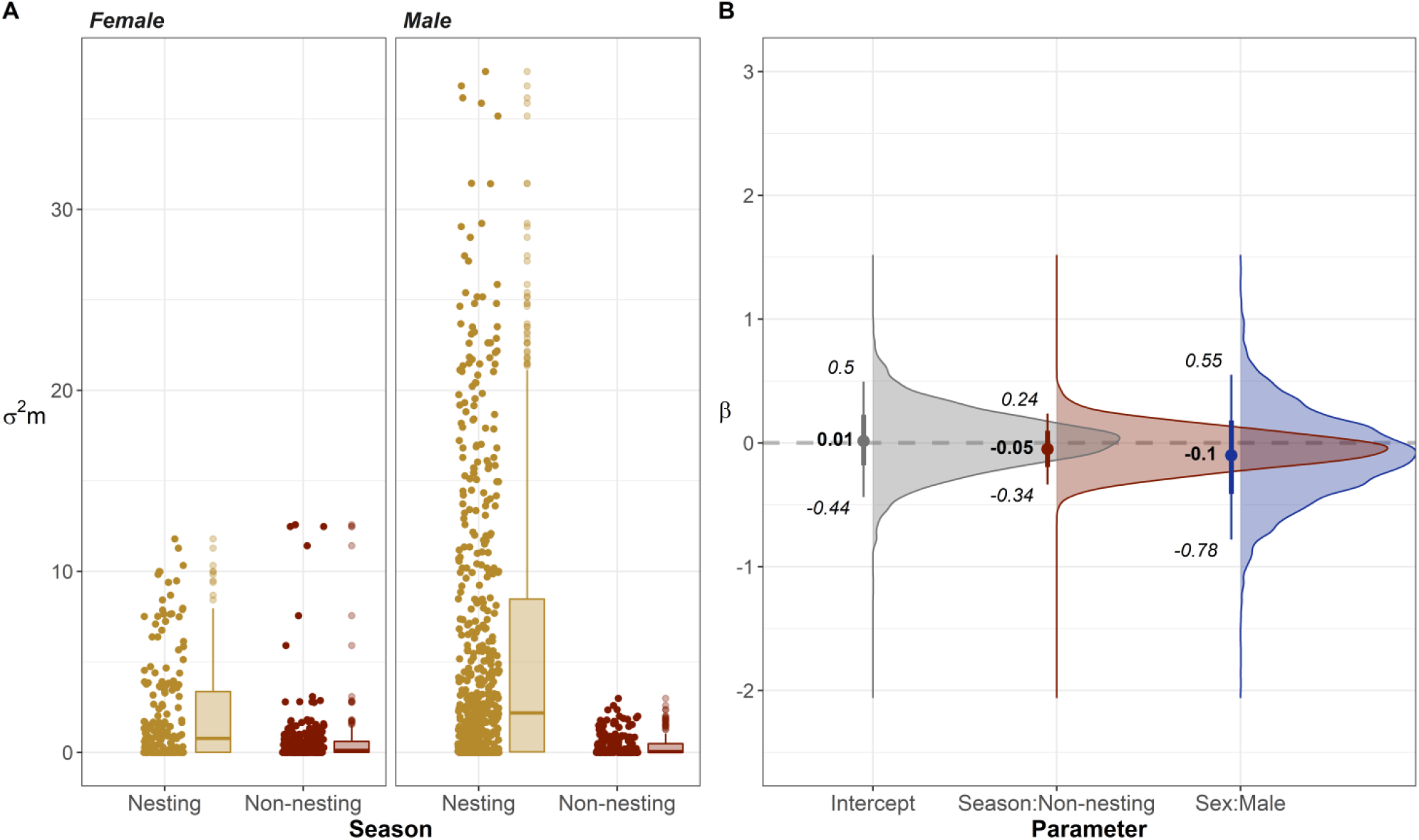
Seasonal differences in motion variance between sexes. **A)** Box and jittered scatter plots of seasonal motion variance values split between female and male. **B)** Model averaging results with point estimates and 95% BCrI.

*Boiga cyanea* depredated avian nests primarily between April and July (Fig. 5). In 2018 and 2019, the nest monitoring cameras recorded fewer nest depredations by *B. cyanea* compared to previous years (Fig. A.5). We attempted to standardize nest depredations by *B. cyanea* accounting for trapping effort: we divided the total number of nests depredated each month by the total number of exposure days per month of the nest monitoring cameras.

**Fig. 5.**
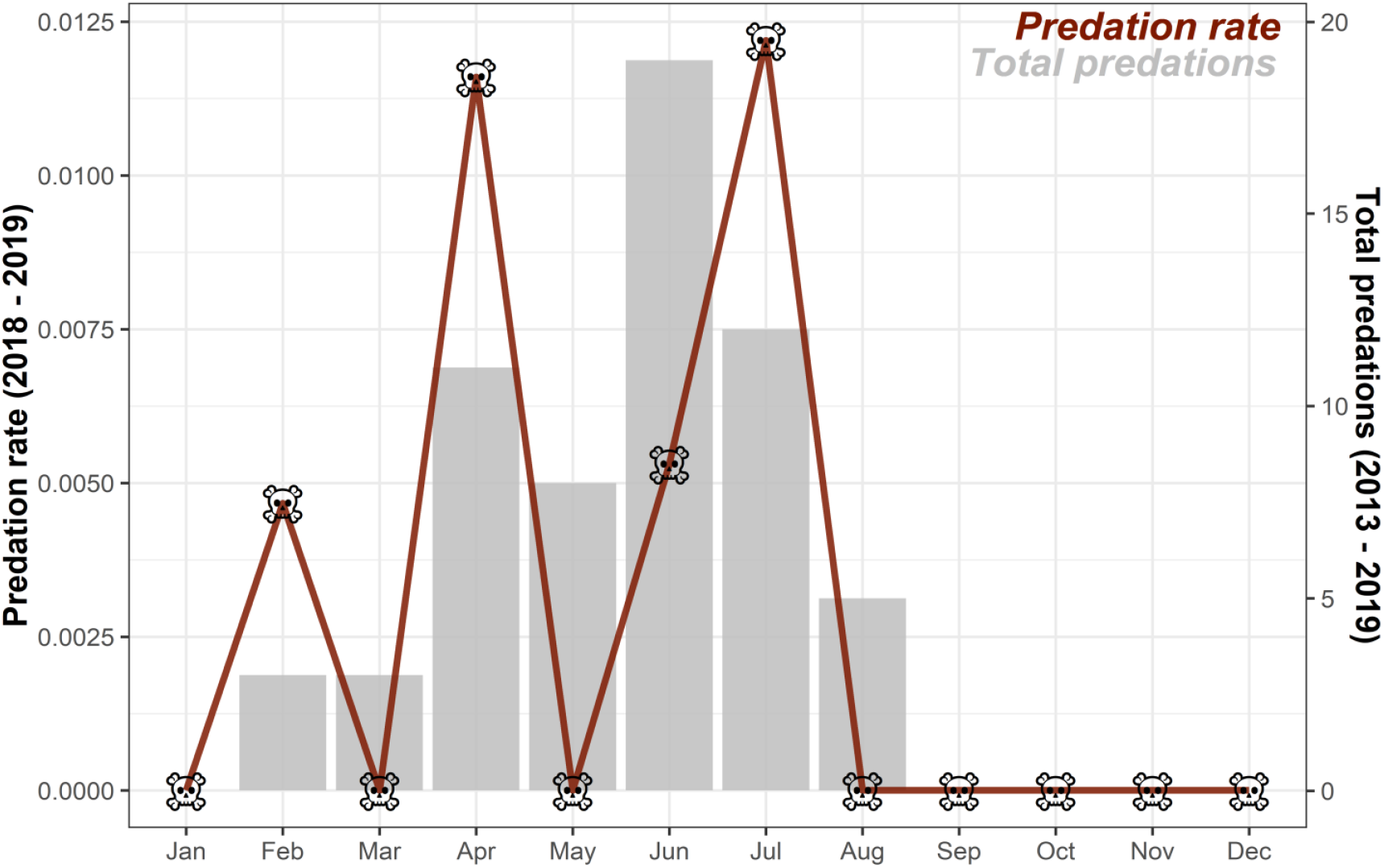
Monthly avian nest depredation rates by *B. cyanea* during the 2018 and 2019 avian nesting seasons, and the total number of nest depredations by *B. cyanea* between 2013 and 2019. These data were obtained via the continuously monitoring nest cameras at SBR.

#### 3.3.2. Nocturnal activity

If snakes had moved from their diurnal refugia by their first nocturnal fix, they generally continued moving throughout the night. We assessed nights with ≥ 2 nocturnal fixes to gauge their activity patterns (Table A.4).

There is considerable temporal overlap (Δ*hat* = 0.68) between *B. cyanea* predation activity recorded by the nest monitoring cameras, and radio-tracked *B. cyanea* foraging activity. Predation activity is concentrated just after dark, while foraging activity appears consistent throughout the night. We observe three snake foraging activity peaks in Fig. 6 as a result of our tracking regime –three night-time fixes spaced apart by approximately four-hour intervals.

**Fig. 6.**
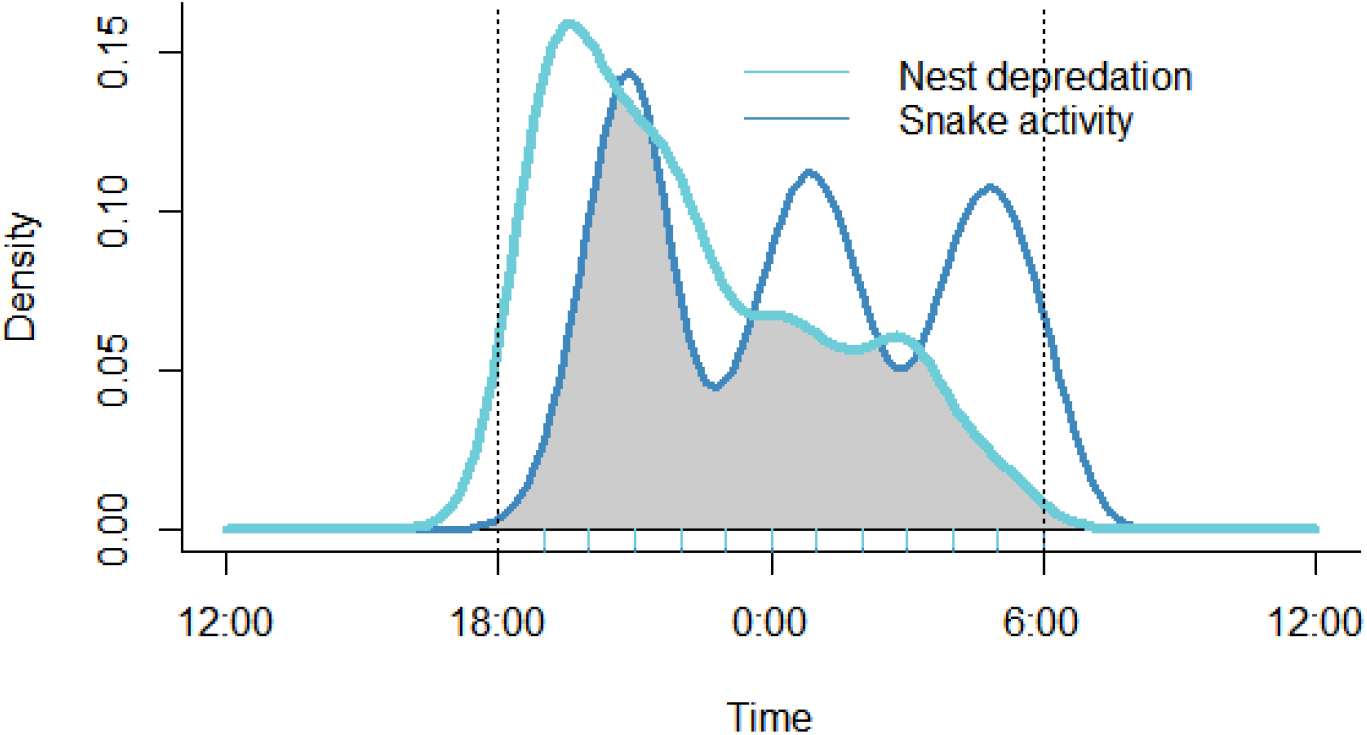
Temporal activity density curves: *B. cyanea* activity (dark blue) and *B. cyanea* nest depredations (light blue). The overlap between the two curves is shaded in grey.

Our snakes generally travelled further at night during the avian nesting season (n = 300) compared to the non-nesting season (n = 110; Fig. 7). Movement and stationary behaviours were significantly different during the avian nesting and non-nesting seasons (Pearson’s Chi-squared Test, p-value = 0.003). Of the 300 nocturnal fixes recorded during the nesting season, snakes were moving 58% of time (n = 174). Of the 110 nocturnal fixes during the non-nesting season, snakes were moving only 41% of the time (n = 45).

**Fig. 7.**
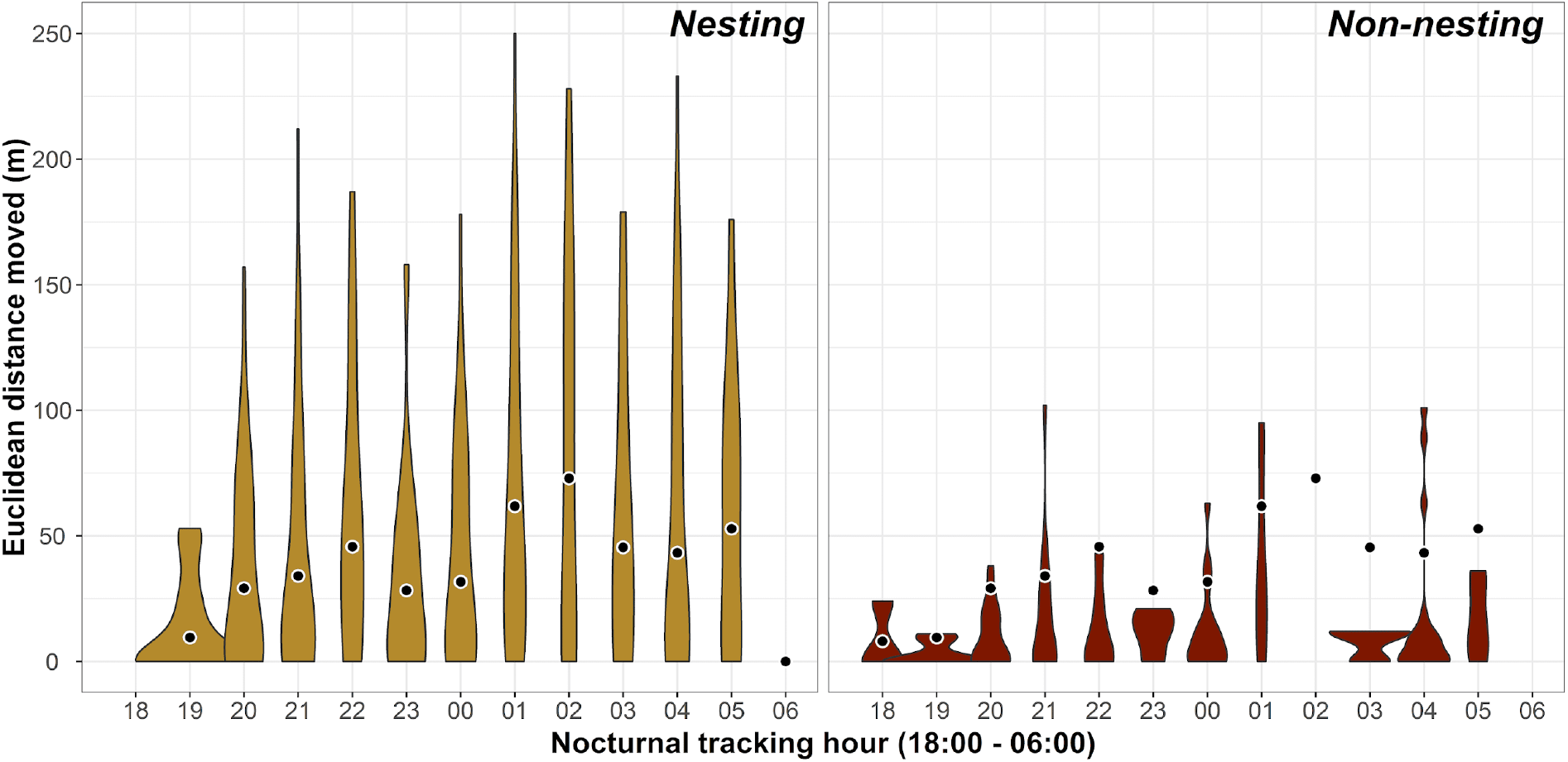
Euclidean distances between snake locations at specific nocturnal hours and previous daytime refugia during the avian nesting season and the non-nesting season. The data illustrated are collected from individuals’ radio-tracked between 2018.03.14 and 2018.09.03, and between 2019.01.15 and 2019.06.08.

## 4. DISCUSSION

Male and female *Boiga cyanea* move and use space in different ways. Our males typically moved two and a half times farther and used areas five and a half times larger during the study. Our male and female *B. cyanea* differed in seasonal activity and space use. During the avian nesting season, radio-tracked *B. cyanea* were approximately seven times more active –moving farther and more frequently (matching our prediction of increased activity during the nesting season). The snakes also used areas six times larger during the avian nesting season compared to the non-nesting season (contrary to our prediction of increased space use during the non-nesting season). Our observations suggest that *B. cyanea* are strictly nocturnal foragers, actively foraging throughout the night. Additionally, our *B. cyanea* typically sheltered at heights above 2 m, and descended closer to the ground to forage at night.

The nesting season –from February to August, partially overlaps with the wet season at the Sakaerat Biosphere Reserve (SBR). The wet season spans from May to October, with typical peaks in May and in September, and low rainfall between June and August. Khamcha *et al*. (2018) reported that rainfall and nest depredations by *B. cyanea* were positively correlated, speculating increased snake activity connected the two. The wet season might trigger a general increase in invertebrate and herpetofauna activity, thereby affecting their interactions with higher trophic levels (Illera and Díaz, 2006; Saenz et al., 2006). With greater prey activity comes greater prey encounter probability. Multiple biotic (e.g. life stage, body condition, prey presence, etc.; Horesh et al., 2017; Kotler et al., 1992) and abiotic factors (e.g. moonlight, temperature, rainfall, etc.; Campbell et al., 2009; Eskew and Todd, 2017) acting synergistically might be driving the increase in *B. cyanea* activity during the avian nesting season. In contrast, Tobin *et al*. (1999) reported that *B. irregularis* on Guam did not exhibit any difference in distances travelled across the rainy and dry seasons. However, Guam as a small island system may have limited seasonal fluctuations in temperature, although situated at similar latitudes to our site. The invasive nature of the snake may inhibit seasonal behaviours, or the system may simply lack seasonal prey availability fluctuations.

Throughout our study, we detected no movement behaviour during daylight hours. Despite non-systematic sampling of diurnal snake behaviour, it appears that *B. cyanea* are active primarily at night. Bulian and Bannasan (1999) report similar ad-hoc observations on *B. cyanea* diel activity, but have observed *B. cyanea* copulating on multiple occasions during the early daylight hours –around 0700 hours. We observed no mating behaviour during our study. Only on one occasion we located our male and female *B*. *cyanea* in the same tree during daylight hours; this could suggest they were possibly copulating but we have no visual evidence of the event. The nest monitoring cameras detected no daytime avian nest depredations from *B. cyanea* further supporting our suspicions of exclusively nocturnal activities (Khamcha et al., 2018).

The tracked individuals often began activity just after sunset, at dusk, and ceased at dawn, just before sunrise. Similar observations exist regarding *Boiga irregularis* nocturnal activity on Guam, with snakes becoming active around sunset, and remaining more or less so throughout the night until ceasing their activity around sunrise (Fritts and Chiszar, 1999; Lardner et al., 2014; Siers et al., 2018). A further similarity with *B. irregularis* may suggest that the stationary periods we observed are the result of prey digestion. On Guam supplementary fed snakes showed lowered activity for 1 to 5 days post feeding (Siers et al., 2018), broadly reflecting our mean stationary period of 3.29 ± 0.66 days.

The snakes were likely to move throughout the night if we recorded moving behaviour by 4 hours post sunset. Thus, we suggest *B. cyanea* are primarily active foragers that might also ambush prey on occasion similar to their invasive congener on Guam, who exhibit a combination of the two foraging modes (Rodda, 1992). We failed to detect any avian nest depredations by radio-telemetered individuals during our study. However, our tracking regime could have allowed our snakes to depredate avian nests during inter-night and intra-night tracking intervals. Low encounter rates of avian nests, active selection of prey sizes, and gape limitations may be reasons why we detected no nest predations during our study (Shine, 1991). Being predominantly active foragers and generalists, *B. cyanea* may still be feeding frequently on other prey (Beaupre and Montgomery, 2007), as SBR hosts a high diversity of sedentary and active prey species (Crane et al., 2018). Our individuals likely fed on smaller but more widespread and abundant prey items, like forest lizards (*Calotes sp.*), geckos or frogs (Bulian and Bannasan, 1999) although frogs in the SBR dry evergreen forest are seasonally restricted to areas adjacent to water.

The nest monitoring cameras detected higher numbers of avian nest depredations during the early hours after sunset, at around 1900 hours. The ornithological researchers observed *B. cyanea* –presumably the same individual, depredating single nest contents from the same nest on consecutive nights, until all the eggs and fledglings were depredated (Personal comm., Angkaew 2017). During these occasions, the predator was likely to have selected refugia relatively close to the avian nests to optimize its feeding benefits while reducing its predation risk. From our radio-tracking data on distances travelled by 1900 hours, ‘relatively close’ would translate to an average distance of 26.2 ± 4.93 m, and a maximum of 53 m. Fledglings and avian eggs might be efficient prey items for *B. cyanea*: the time taken to subdue and ingest these prey might be greatly reduced compared to larger and effectively defensive prey (Pleguezuelos et al., 2007), thus reducing the predator’s own predation risk (Mullin and Cooper, 1998).

The tracked snakes sheltered at heights over 2 m off the ground during most diurnal fixes (≈95%). Thus *B. cyanea* must frequently descend from their diurnal refugia to lower heights at night to forage. Our findings are consistent with descriptions of the Dark-headed cat snake *Boiga nigriceps* (Fujishima et al., 2021) and *B. irregularis* (Rodda, 1992) foraging heights. The nest predation records show that White-rumped Shamas *Kittacincla malabarica* (WRSH), Abbott’s babbler *Malacocincla abbotti* (ABBA), and Scaly-crowned Babblers *Malacopteron cinereum* (SCBA) were the three most depredated species’ nests by *B. cyanea* at SBR. Khamcha and Gale (2020) report *B. cyanea* being responsible for ≈27% of predations by top-five predators on WRSH, ≈25% on ABBA, and ≈19% on SCBA. The average nest height for WRSH is around 1.6 m (Khamcha and Gale, 2020), that for ABBA is 0.7 m (Khamcha and Gale, 2020), and that for SCBA is 0.98 m (Somsiri et al., 2020) –similar to the median movement height (1.5 m; IQR: 3.5) recorded for our *B. cyanea*. *Boiga cyanea* might forage closer to the ground, especially during the avian nesting season, to actively hunt for ground and understory nesting bird nests.

Our study is the first to explore the space use and activity patterns of a regionally important snake predator in relation to its prey. We attempted to quantify *B. cyanea* foraging activity by implicitly assuming their movements, particularly during the avian nesting season, were related to foraging rather than other biological functions. For this reason, our findings regarding *B. cyanea* foraging behaviour must not be overstated. We developed our nocturnal tracking protocol to minimize our influence on the snakes’ natural behaviours; locating each individual every night, rather than alternate nights, would have provided more detailed insights into their nocturnal activity patterns while potentially revealing movement patterns occurring at different temporal scales. The disparity in total fixes between the avian nesting and non-nesting seasons hindered us from making robust statistical inferences on seasonal horizontal and vertical movements. Having no radio-tracking overlap across the seasons for most individuals (n = 9) further limits our inferences on seasonal changes in activity patterns.

We caution against extrapolating seasonal space use and activity patterns of males and females, because of possible sources of sampling biases, as detailed in the STRANGE framework (Webster and Rutz, 2020). **Social background**: individuals’ size might have influenced their movements with respect to their conspecifics, either through avoidance or attraction behaviour. **Trappability and self-selection:** individuals predisposed to moving towards anthropogenic habitat features may have been more likely to be captured. Because most individuals were captured opportunistically (n = 9), our sample is non-random and non-representative of the *B. cyanea* population at large. **Acclimation and habituation:** our presence during nighttime fixes likely caused behavioural changes despite our efforts to minimize disturbance. **Natural changes in responsiveness:** the avian nesting seasons in 2018 and 2019 witnessed lower nest depredations by *B. cyanea* compared to 2013 – 2017. Reasons for the two anomalous years are unknown. The association between *B. cyanea* activity and nest depredations during our study period, might therefore not reflect that of previous years, or of those to come.

Our study highlights the need to explore *B. cyanea* diet to quantify what proportion fledglings and avian eggs constitute at SBR. Studying patterns of prey selection (whether active or opportunistic) and understanding how *B. cyanea* locate avian nests could help better elucidate these complex predator-prey relationships. Bird populations at SBR face significant predation pressure from both nocturnal snakes and the Northern Pig-tailed Macaque *Macaca leonina* (Kaisin et al., 2018; Khamcha et al., 2018). Understanding the combined effect of these major sympatric predators on nest survival could help develop strategies to increase nest success. The results of this study might be applicable to other parts of Southeast Asia where snakes might be important avian nest predators, especially of bird populations of conservation concern (e.g. Donald et al., 2009).

## DECLARATION OF COMPETING INTEREST

The authors declare that they have no known competing financial interests or personal relationships that could have appeared to influence the work reported in this paper.

## ACKNOWLEDGEMENTS

We sincerely thank the National Science and Technology Development Agency of Thailand (NSTDA) for funding this research project (grant number: P-17-50476). We thank the National Research Council of Thailand (NRCT) for issuing our research permit (Number: 0002/4493). We thank the Institute of Animals for Scientific Purpose Development (IAD) for issuing our animal use license (Number: 1-08385-2562). We thank the Thailand Institute of Scientific and Technological Research for granting permission to conduct research within the Sakaerat Biosphere Reserve (SBR). We sincerely thank the Sakaerat Environmental Research Station (SERS) for logistic support throughout the duration of the project. We thank the Suranaree University of Technology for bureaucratic assistance. We thank Nakhon Ratchasima zoo, in particular D.V.M. Wirongrong Changphet, for providing veterinary expertise. We thank all fellow researchers, interns and staff at SERS for their contributions towards the running of this project. Lastly, a very special thanks to field technicians Nathaniel Quarrell, Jizel Miles, Thomas Prewett, Rose Stroup and Harry Ward-Smith for their time, devotion, enthusiasm, hard work and work ethic.

## Funding

This work was supported by the National Science and Technology Development Agency of Thailand (grant number: P-17-50476, 2017).

## AUTHOR CONTRIBUTIONS

Conceptualization: A.D., C.T.S., G.G., and D. K.; Methodology: A.D., C.T.S., G.G., and B.M.M.; Formal Analysis: A.D. and B.M.M.; Investigation: A.D.; Resources: A.D., C.T.S., G.G., D. K., and S.W.; Writing – Original Draft: A.D. and C.T.S.; Writing – Review & Editing: A.D., C.T.S., G.G., B.M.M., D. K., and S.W.; Visualization: A.D. and B.M.M.; Supervision: C.T.S., G.G., and S.W.; Funding Acquisition: A.D., C.T.S., G.G., D. K., and S.W. The author(s) read and approved the final manuscript.

## APPENDIX A.

**Fig. A.1.**
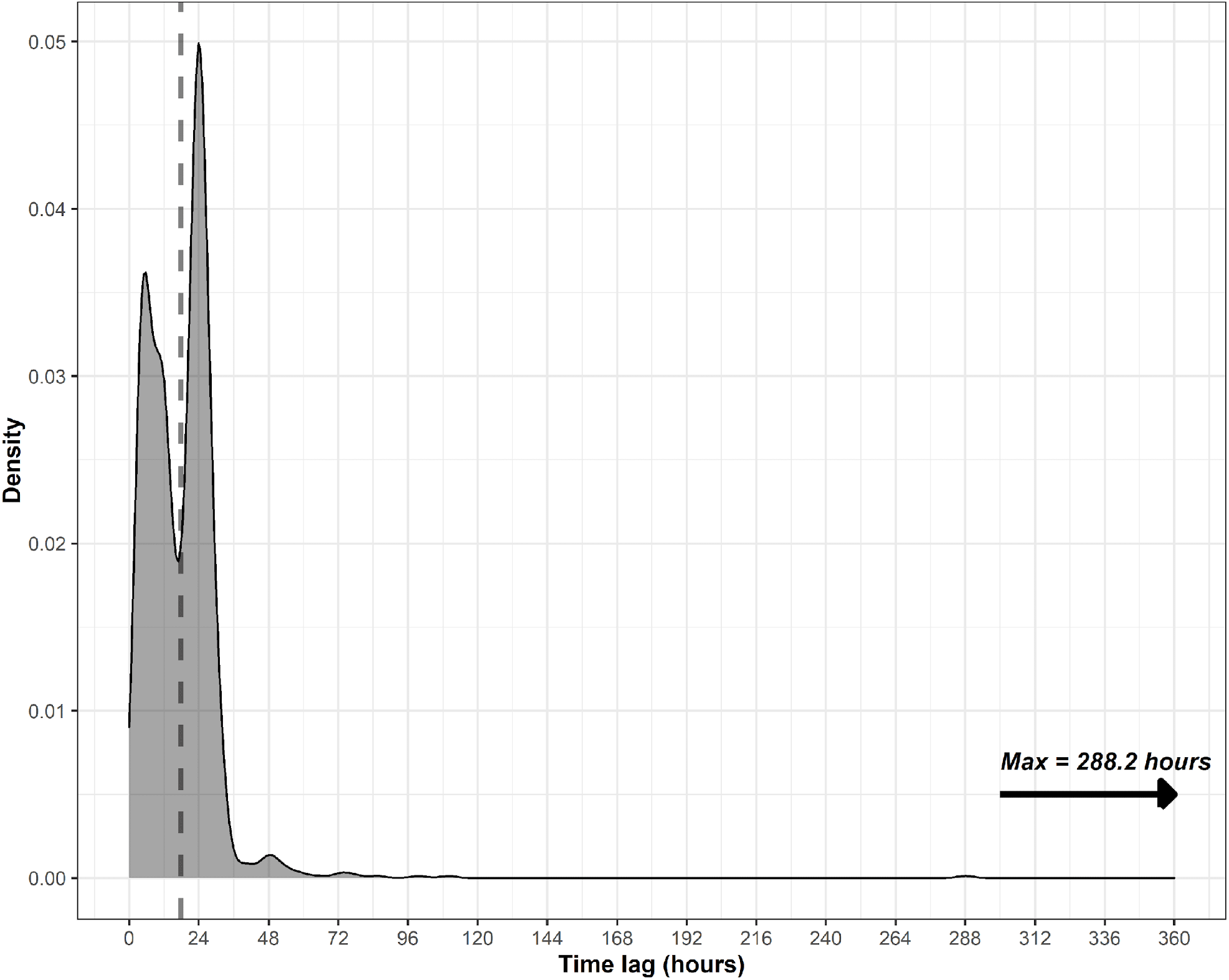
Average tracking interval (17.63 ± 0.37 hours) between diurnal and nocturnal fixes of our radio-tracked individuals. The peak around 48 hours is likely the result of times we were unable to detect radio-signal on our individuals. The peaks between 72 hours and 288 hours are primarily the result of snakes were awaiting transmitter replacement surgery at the research facility.

**Fig. A.2.**
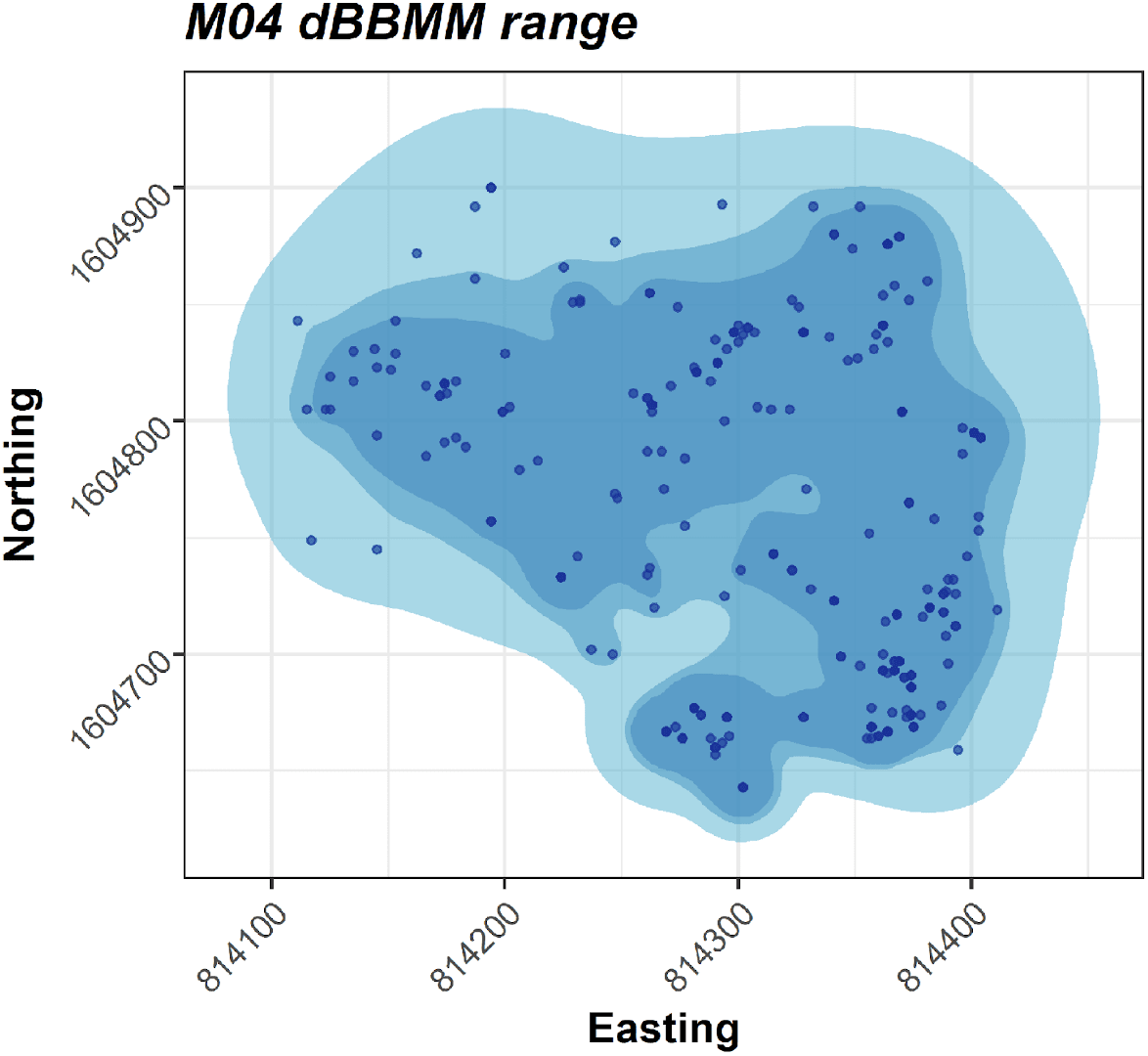
99%, 95%, and 90% dBBMM isopleth contours M04, radio-tracked 6 November 2018 and 8 June 2019.

**Fig. A.3.**
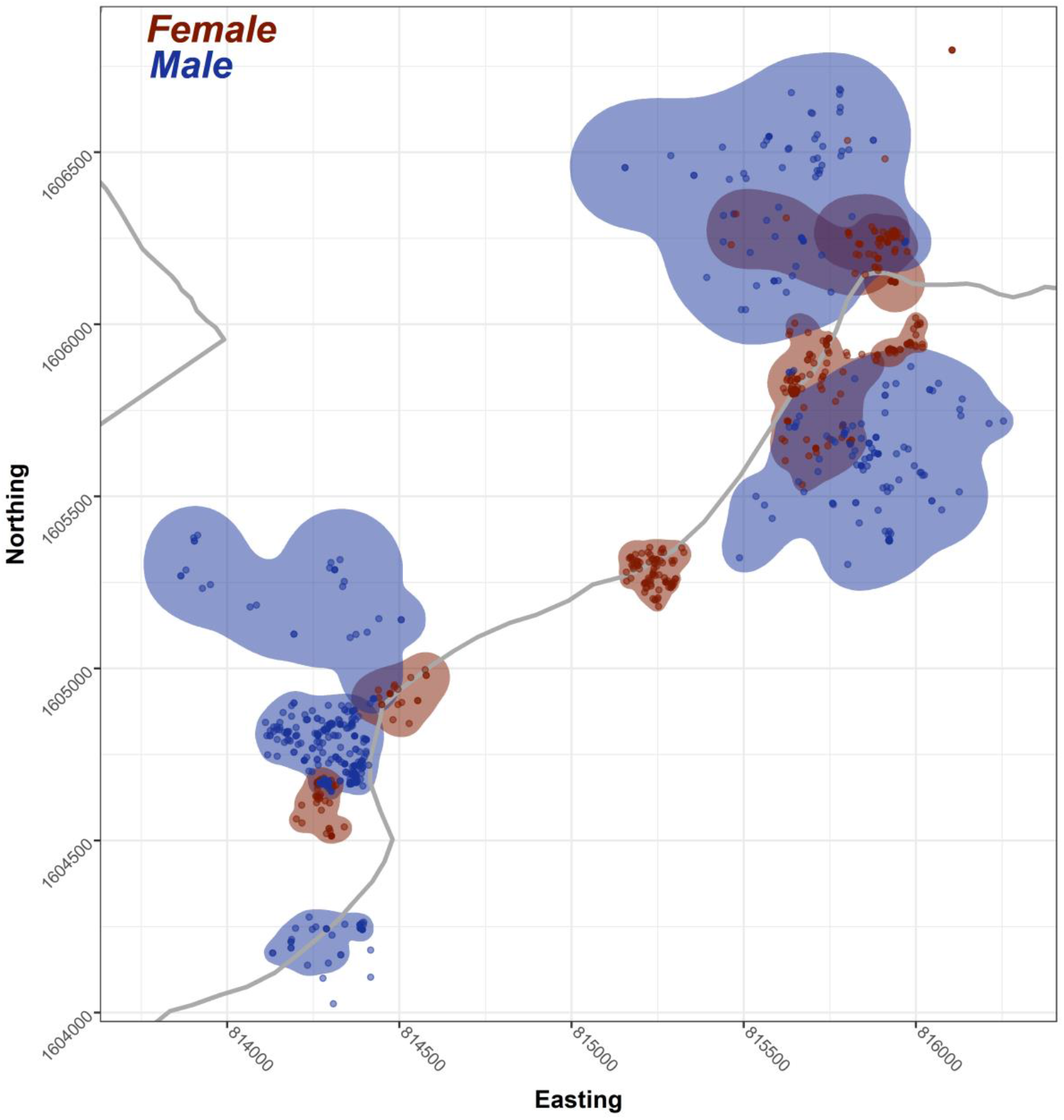
DBBMM occurrence distributions illustrating 99% isopleth contours for male (blue) and female (red) radio-tracked *B. cyanea* between 21 October 2017 and 8 June 2019 in reference to the main roads (grey) within the SBR core area.

**Fig. A.4.**
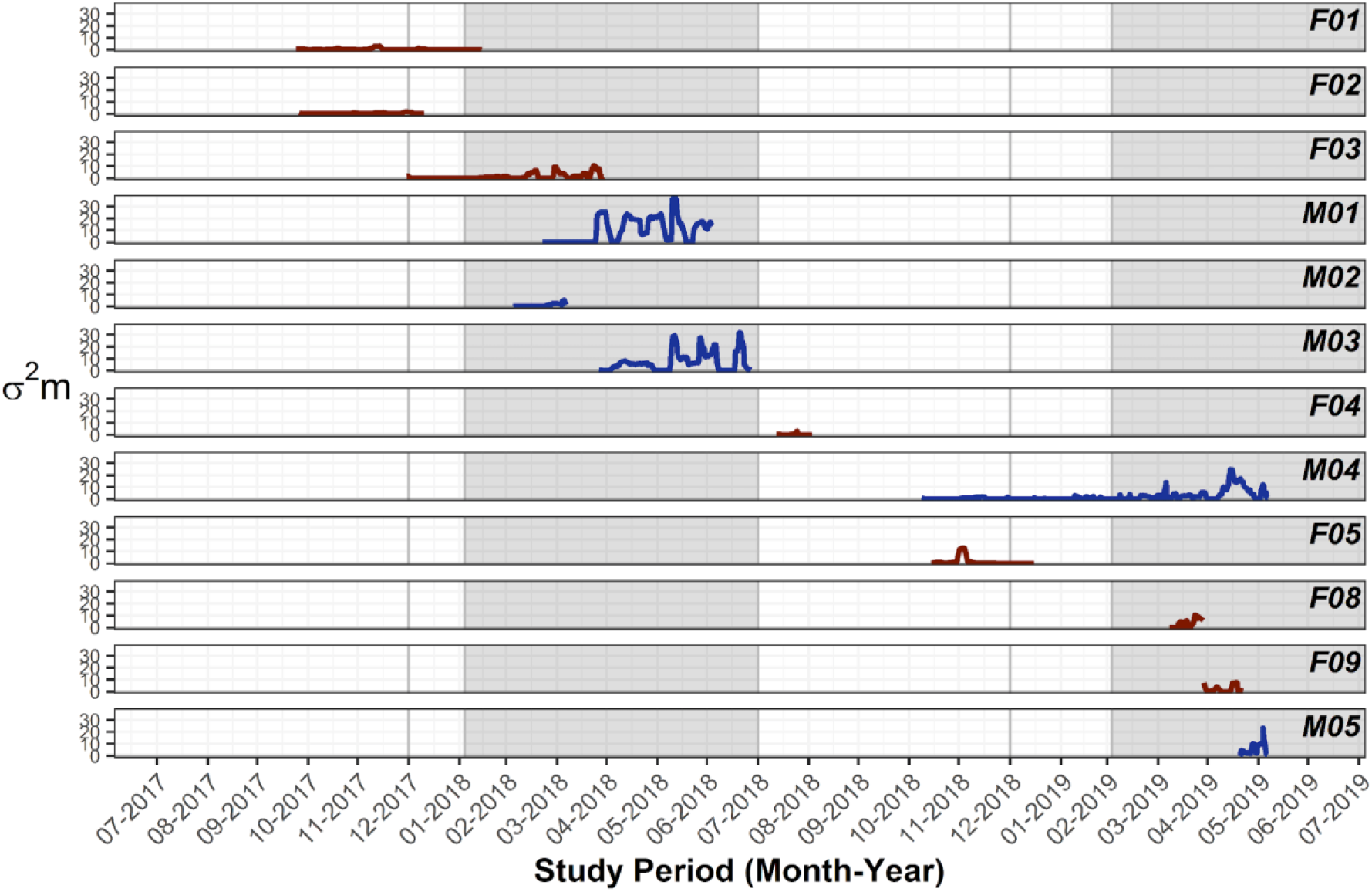
Individual motion variances for radio-tracked *B. cyanea* with nesting seasons 2018 and 2019 evidenced in grey.

**Fig. A.5.**
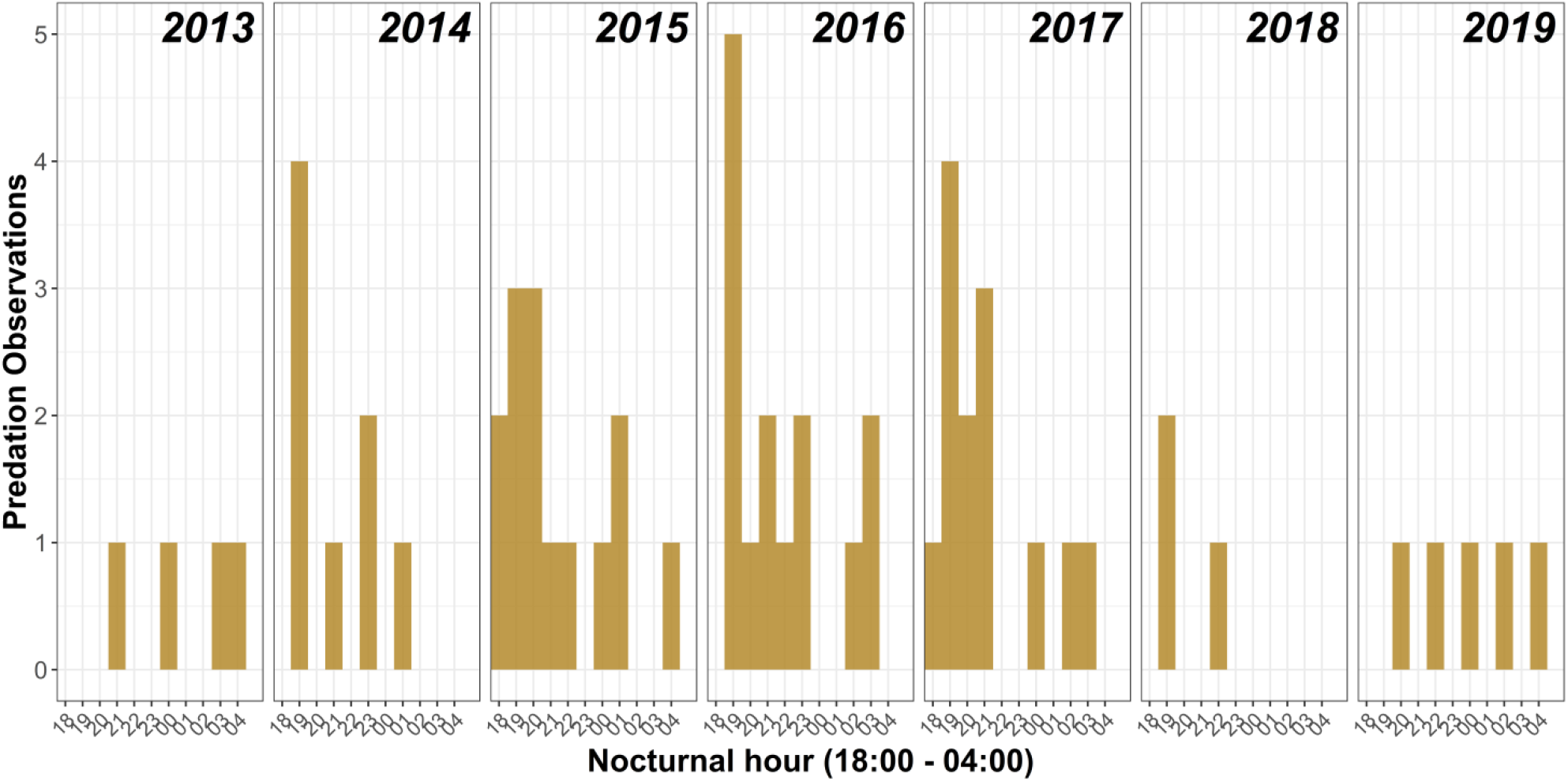
Avian nest depredations by *B. cyanea* broken down by hour and by year, recorded via the continuously monitoring nest cameras at SBR.

**Fig. A.6.**
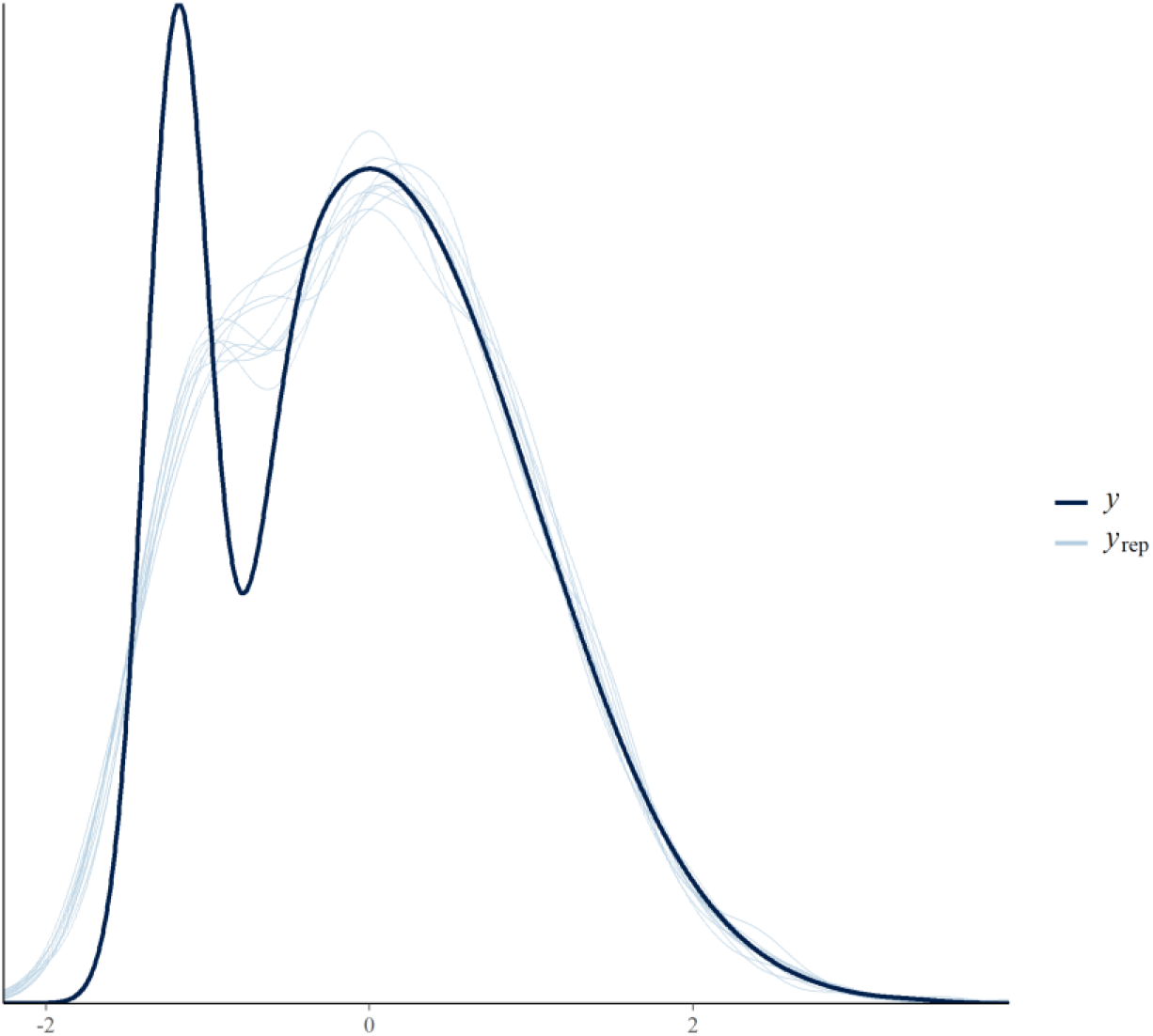
Posterior predictive check for Bayesian regression model plotting observed data (y) and simulated data (y_rep_).

**Table A.1.** Occurrence distributions and motion variances of radio-tracked *Boiga cyanea* between 21 October 2017 and 8 June 2019 using window size 11 and margin size 5.

| Snake ID | Days Tracked | 90% dBBMM (ha) | 95% dBBMM (ha) | 99% dBBMM (ha) | $\sigma_m^2 \pm SE$ |
| --- | --- | --- | --- | --- | --- |
| F01 | 119 | 1.63 | 2.10 | 3.11 | $0.53 \pm 0.06$ |
| F02 | 82 | 0.98 | 1.53 | 2.64 | $0.37 \pm 0.05$ |
| F03 | 124 | 4.17 | 6.40 | 11.25 | $1.52 \pm 0.19$ |
| F04 | 24 | 0.56 | 0.82 | 1.38 | $0.34 \pm 0.1$ |
| F05 | 65 | 1.58 | 4.88 | 12.53 | $1.27 \pm 0.35$ |
| F08 | 22 | 2.24 | 3.26 | 5.72 | $1.33 \pm 0.37$ |
| F09 | 27 | 1.88 | 2.70 | 4.72 | $2.23 \pm 0.45$ |
| M01 | 105 | 28.13 | 37.54 | 57.78 | $8.59 \pm 0.91$ |
| M02 | 36 | 1.90 | 2.73 | 4.06 | $0.66 \pm 0.14$ |
| M03 | 95 | 18.43 | 24.88 | 39.89 | $8.02 \pm 0.78$ |
| M04 | 214 | 3.93 | 5.21 | 8.58 | $2.17 \pm 0.21$ |
| M05 | 19 | 16.70 | 21.08 | 30.61 | $7.44 \pm 1$ |

\*dBBMM: Dynamic Brownian bridge movement model estimate; \* $\sigma_m^2 \pm SE$ : Motion variance and standard error.

**Table A.2.** Occurrence distributions and motion variances of radio-tracked *Boiga cyanea* between 21 October 2017 and 8 June 2019 using window size 15 and margin size 7.

| Snake ID | Days Tracked | 90% dBBMM (ha) | 95% dBBMM (ha) | 99% dBBMM (ha) | $\sigma_m^2 \pm SE$ |
| --- | --- | --- | --- | --- | --- |
| F01 | 119 | 1.70 | 2.17 | 3.15 | $0.5 \pm 0.05$ |
| F02 | 82 | 0.99 | 1.48 | 2.45 | $0.35 \pm 0.05$ |
| F03 | 124 | 3.86 | 5.89 | 10.27 | $1.64 \pm 0.21$ |
| F04 | 24 | 0.53 | 0.77 | 1.29 | $0.32 \pm 0.09$ |
| F05 | 65 | 2.19 | 5.53 | 12.68 | $1.33 \pm 0.33$ |
| F08 | 22 | 1.88 | 2.60 | 4.26 | $1.31 \pm 0.23$ |
| F09 | 27 | 1.81 | 2.52 | 4.19 | $2.11 \pm 0.41$ |
| M01 | 105 | 27.68 | 36.31 | 54.35 | $8.37 \pm 0.81$ |
| M02 | 36 | 1.80 | 2.67 | 4.16 | $0.69 \pm 0.16$ |
| M03 | 95 | 18.70 | 25.06 | 39.66 | $8.11 \pm 0.67$ |
| M04 | 214 | 4.01 | 5.27 | 8.57 | $2.08 \pm 0.19$ |
| M05 | 19 | 11.86 | 15.34 | 22.92 | $7.85 \pm 0.88$ |

\*dBBMM: Dynamic Brownian bridge movement model estimate; \* $\sigma_m^2 \pm SE$ : Motion variance and standard error.

**Table A.3.**
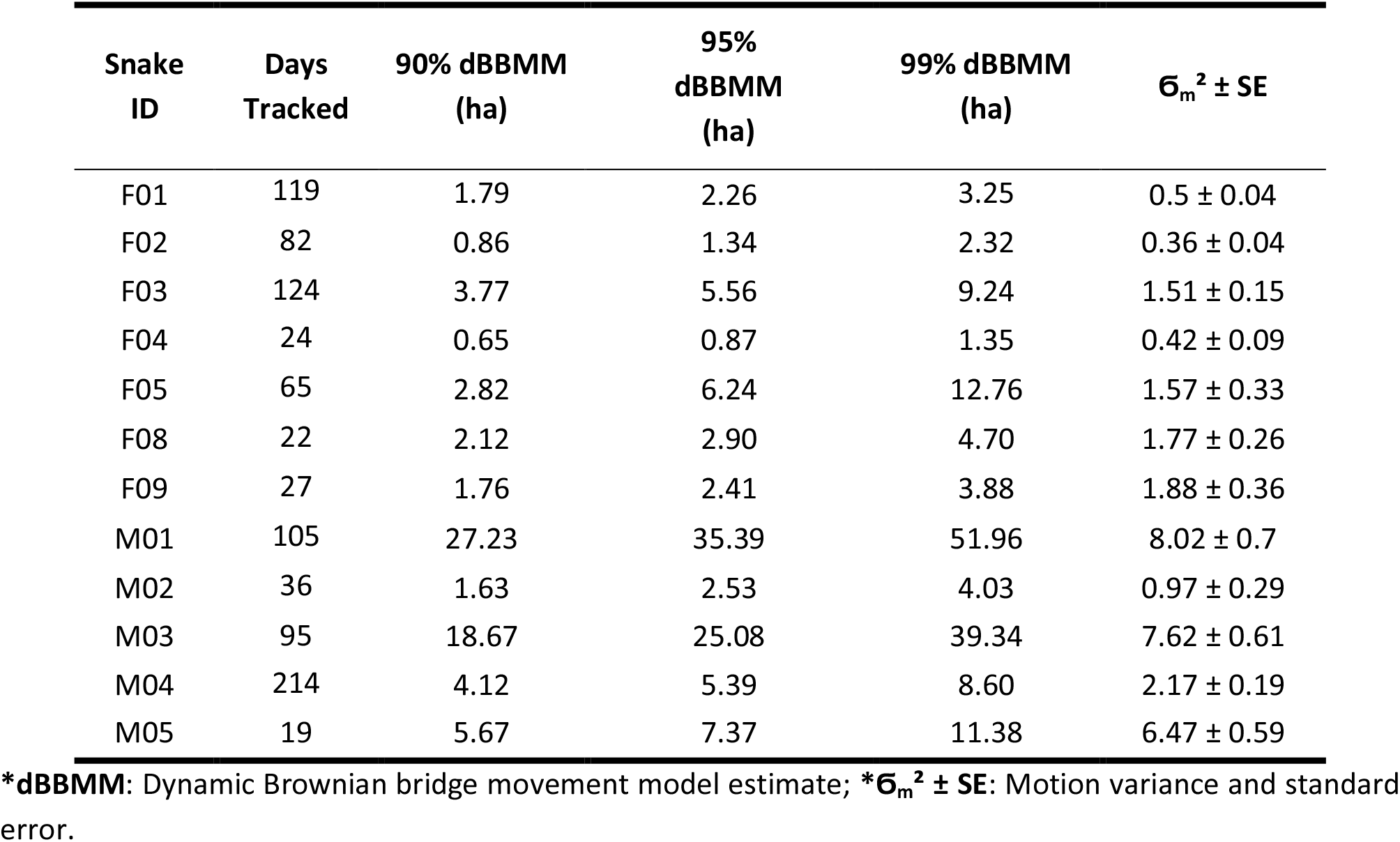
Occurrence distributions and motion variances of radio-tracked *Boiga cyanea* between 21 October 2017 and 8 June 2019 using window size 21 and margin size 9.

| Snake ID | Days Tracked | 90% dBBMM (ha) | 95% dBBMM (ha) | 99% dBBMM (ha) | $\sigma_m^2 \pm SE$ |
| --- | --- | --- | --- | --- | --- |
| F01 | 119 | 1.79 | 2.26 | 3.25 | $0.5 \pm 0.04$ |
| F02 | 82 | 0.86 | 1.34 | 2.32 | $0.36 \pm 0.04$ |
| F03 | 124 | 3.77 | 5.56 | 9.24 | $1.51 \pm 0.15$ |
| F04 | 24 | 0.65 | 0.87 | 1.35 | $0.42 \pm 0.09$ |
| F05 | 65 | 2.82 | 6.24 | 12.76 | $1.57 \pm 0.33$ |
| F08 | 22 | 2.12 | 2.90 | 4.70 | $1.77 \pm 0.26$ |
| F09 | 27 | 1.76 | 2.41 | 3.88 | $1.88 \pm 0.36$ |
| M01 | 105 | 27.23 | 35.39 | 51.96 | $8.02 \pm 0.7$ |
| M02 | 36 | 1.63 | 2.53 | 4.03 | $0.97 \pm 0.29$ |
| M03 | 95 | 18.67 | 25.08 | 39.34 | $7.62 \pm 0.61$ |
| M04 | 214 | 4.12 | 5.39 | 8.60 | $2.17 \pm 0.19$ |
| M05 | 19 | 5.67 | 7.37 | 11.38 | $6.47 \pm 0.59$ |

\*dBBMM: Dynamic Brownian bridge movement model estimate; \* $\sigma_m^2 \pm SE$ : Motion variance and standard error.

**Table A.4.** Tracking interval summaries of nights with ≥ 2 nocturnal fixes, during which the snakes had moved or were observed moving.

| Nocturnal fix | Fixes | Moved | Moving |
| --- | --- | --- | --- |
| 1<sup>st</sup> interval (18:00-21:59) | 110 (35%) | 66 (34%) | 62 (37%) |
| 2<sup>nd</sup> interval (22:00-01:59) | 97 (31%) | 64 (33%) | 58 (35%) |
| 3<sup>rd</sup> interval (02:00-07:00) | 105 (34%) | 66 (34%) | 47 (28%) |
| **Total** | 312 | 196 (63%) | 197 (54%) |

## Notes

https://osf.io/6yrbg/

### Competing Interest Statement

The authors have declared no competing interest.

